# Terminal uridylyltransferases target RNA viruses as part of the innate immune system in animals

**DOI:** 10.1101/209114

**Authors:** Jérémie Le Pen, Hongbing Jiang, Tomás Di Domenico, Emma Kneuss, Joanna Kosałka, Marcos Morgan, Christian Much, Konrad L. M. Rudolph, Anton J. Enright, Dónal O’Carroll, David Wang, Eric A. Miska

## Abstract

RNA viruses are a major threat to animals and plants. RNA interference (RNAi) and the interferon response provide innate antiviral defense against RNA viruses. Here we performed a large-scale screen using *C. elegans* and its natural pathogen, the Orsay virus (OrV), and identified *cde-1* as important for antiviral defense. CDE-1 is a homologue of the mammalian TUT4/7 terminal uridylyltransferases; its catalytic activity is required for its antiviral function. CDE-1 uridylates the 3′ end of the OrV RNA genome and promotes its degradation, independently of the RNAi pathway. Likewise, TUT4/7 uridylate influenza A virus (IAV) mRNAs in mammalian cells. Deletion of TUT4/7 leads to increased IAV mRNA and protein levels. We have defined 3′ terminal uridylation of viral RNAs as a conserved antiviral defense mechanism.

## MAIN

RNA viruses are a major threat to human health and food security. RNA interference (RNAi) is an important antiviral pathway in most animals and plants: Dicer recognizes and cleaves the double-stranded viral RNA genome into virus-derived small interfering RNAs (viral siRNAs, viRNAs), which are loaded into Argonaute proteins to form the RNA-induced silencing complex (RISC) that in turn targets the viral RNA genome ^1^. Vertebrates have additionally evolved a cellular signaling-based pathway, the interferon response (IR): upon recognition of foreign RNAs (i.e. double-stranded or bearing a 5′ di/triphosphate), cytosolic receptors of the RIG-I family activate the IR which results in an antiviral state of the cell ^2,3^. Here, we identify 3′ terminal uridylation of viral RNAs as a third antiviral mechanism in animals (Extended Data Fig. 1).

We carried out a forward genetic screen to discover antiviral pathways in animals using the nematode *Caenorhabditis elegans* and its natural intestinal pathogen, the Orsay virus (OrV) ^4-12^ (Extended Data Fig. 1). While *C. elegans* lacks an interferon pathway, a RIG-I ortholog, DRH-1, acts in viral recognition. DRH-1 forms a Viral Recognition Complex (ViRC) with the *C. elegans* Dicer (DCR-1) and the RNA-binding protein RDE-4 to link viral recognition to a dedicated antiviral RNAi pathway ^5,11,13,14^ (Extended Data Fig. 1). DRH-1 also induces a transcriptional immune response through an as yet uncharacterized signaling pathway ^10,15^. We generated a viral stress sensor transgene by placing the green fluorescent protein (GFP) under the control of the *lys-3* promoter (allele *mjIs228),* an immune response gene (Fig. 1a and Extended Data Fig. 2a). Upon infection, the level of GFP expression in the intestine mirrored the viral load in wild type, *drh-1* and *rde-1* mutants (Extended Data Fig. 2b, c). We used chemical mutagenesis to screen ~50,000 haploid genomes (Fig. 1b) and identified 16 isolates we named Ovid (Orsay Virus Immune Deficient; Fig. 1c and Extended Data Table 1). 13 out of 16 *ovid* mutants showed increased viral loads (Fig. 1c). *ovid-3,4,5,10,12* are compromised in somatic RNAi (Fig. 1c) and *ovid-3,4,10* carry new alleles of RNAi genes *mut-16, rde-4* and *rrf-1,* respectively (Table 1). To further stratify our Ovid isolates, we assayed DRH-1 pathway activation using the expression of a downstream induced gene named *sdz-6* as readout ^10^ (Fig. 1d and Extended Data Fig. 2a). Only *ovid-1* phenocopied *drh-1* mutants, defining a new allele of *drh-1* (Fig. 1d). We identified a number of additional candidate genes (Table 1). *ovid-9* and *ovid-11* mutants are neither defective in canonical RNAi nor in the DRH-1 pathway and thus represent candidate genes for novel antiviral defense mechanisms.

**Table 1.** Ovid screen candidate genes

| Genotype | High viral load? | RNAi intact? | High sdz-6 level? | Candidate gene | Candidate variation | Brief description |
| --- | --- | --- | --- | --- | --- | --- |
| WT | No | Yes | Yes |  |  |  |
| *rde-1* | Yes | No | Yes |  |  | RNAi factor |
| *drh-1* | Yes | Yes | No |  |  | Viral RNA receptor |
| *ovid-1* | Yes | Yes | No | *drh-1* | Glu834Lys | Viral RNA receptor |
| *ovid-2* | Yes | Yes | Yes | n.d. |  |  |
| *ovid-3* | Yes | No | Yes | *mut-16* | Gln861* | RNAi factor |
| *ovid-4* | Yes | No | Yes | *rde-4* | Ala220Thr | RNAi factor |
| *ovid-5* | Yes | No | Yes | n.d. |  |  |
| *ovid-6* | Yes | Yes | Yes | T09B4.2 | Pro330Leu | Putative rho guanine nucleotide exchange factor |
| *ovid-7* | Yes | Yes | Yes | C41D11.6 | Gly596Ser | Putative RNA nuclease |
| *ovid-8* | Yes | Yes | Yes | n.d. |  |  |
| *ovid-9* | Yes | Yes | Yes | *cde-1* | Gln910* | Terminal uridylyltransferase |
| *ovid-10* | Yes | No | Yes | *rrf-1* | Gly45Glu | RNAi factor |
| *ovid-11* | Yes | Yes | Yes | C54D10.14 | Gly122Arg | Uncharacterized, DRH-1-dependent induction |
| *ovid-12* | Yes | No | Yes | F27D4.6 | Arg717* | Uncharacterized |
| *ovid-13* | n.s. | Yes | Yes | n.d. |  |  |
| *ovid-14* | n.s. | Yes | Yes | n.d. |  |  |
| *ovid-15* | n.s. | Yes | Yes | n.d. |  |  |
| *ovid-16* | Yes | Yes | Yes | *phi-32*<br>*ssl-1* | Pro75Ser<br>Gly1119Glu | Ubiquitin gene<br>SNF2-related |

n.s., not scored. n.d., not determined.

**Figure 1.**
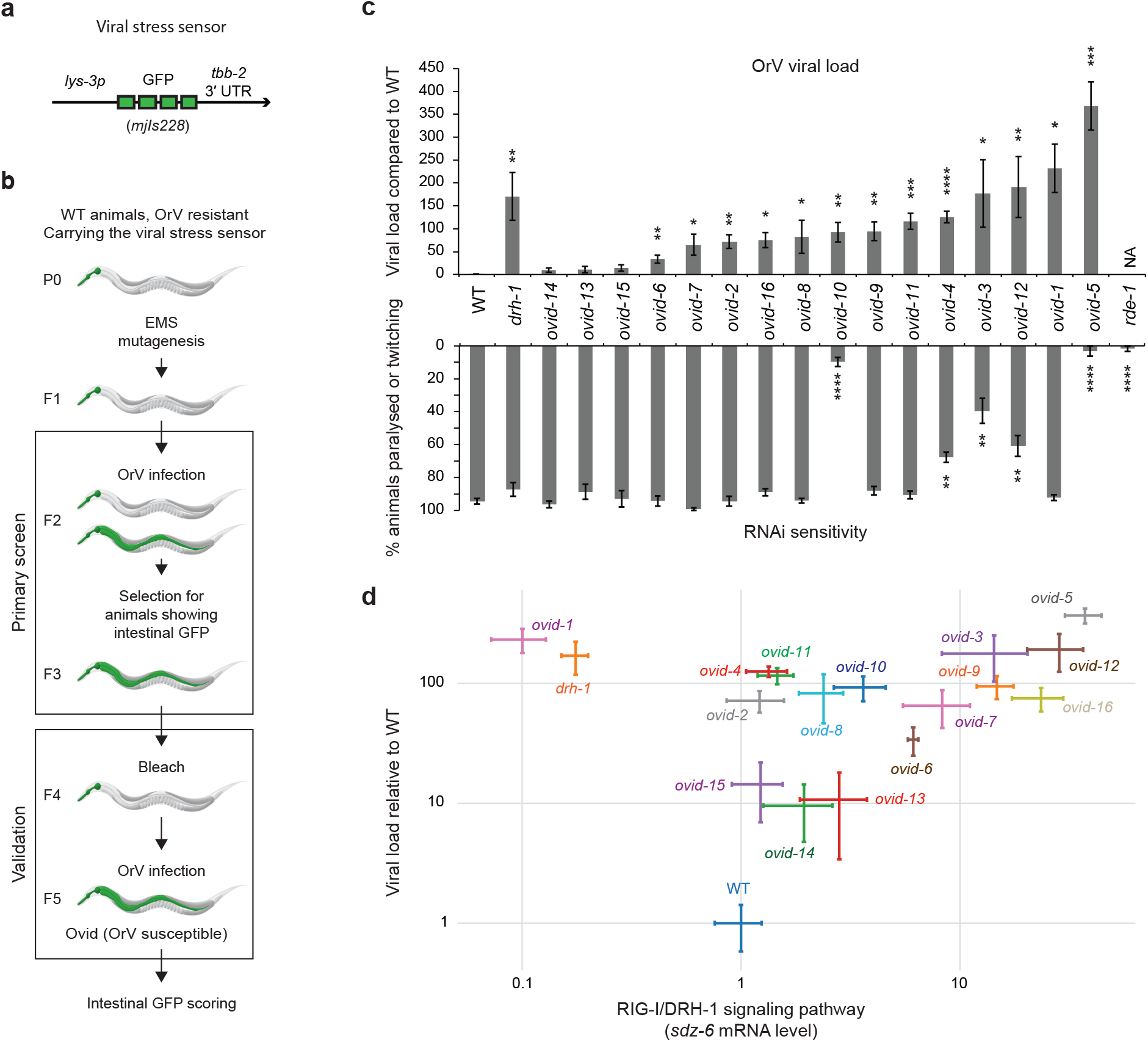
A forward genetic screen identifies novel antiviral immunity genes. **a**, Diagram of the *lys-3p::gfp* viral stress sensor. **b**, Ovid screen workflow. Transgenic animals carrying the viral stress sensor were mutagenized using EMS and F2 progeny were assayed. OrV, Orsay virus. Ovid, Orsay virus immunodeficient. **c**, Top panel: viral load of strains as indicated, measured by qRT-PCR of OrV RNA1, 4 dpi. Error bars represent the standard error of the mean (SEM) of four biological replicates. Onetailed student’s t-test: ****p<0.0001, *** p<0.001, **p<0.01, *p<0.05. Bottom panel: locomotion defects scored (paralyzed or twitching) after *unc-22* RNAi feeding. Error bars: SEM, three biological replicates. Two-tailed student’s t-test: ****p<0.0001, **p<0.01. **d**, Viral load compared to *sdz-6* mRNA levels by qRT-PCR. Error bars: SEM of four biological replicates. Samples as in **c**.

Whole-genome re-sequencing and genetic complementation tests revealed the causative mutation in *ovid-9* to be a single-nucleotide nonsense mutation in the *cde-1* gene *(mj414,* glutamine 910 to STOP) (Fig. 2a and Extended Data Fig. 3). *cde-1* encodes a catalytically active 3′-terminal RNA uridylyltransferase (TUT), which is a homologue of mammalian TUT4 and TUT7 enzymes ^16-18^ (Fig. 2a). The independently derived *cde-1 (tm1021)* knockout strain also phenocopied viral stress sensor activation (Extended Data Fig. 4), high viral loads (Fig. 2b), and horizontal transmission of infection (Extended Data Fig. 4). RNA FISH revealed that viral infection is restricted to the intestine in *cde-1* and in *cde-1; drh-1* double mutants ^4,9^ (Extended Data Fig. 5a). We validated that CDE-1 is present in the intestine using a GFP fusion ^17^ (Extended Data Fig. 5b). To disentangle between the functions of CDE-1 in different tissues, *cde-1* was exclusively expressed from an intestine-specific *vha-6p* promoter (Extended Data Fig. 5c). Animals with intestinal expression of *cde-1* became resistant to viral infection (Extended Data Fig. 5d), but kept a defect in meiotic chromosome segregation (Extended Data Fig. 5e), probably caused by CDE-1 depletion in the germline ^16^. CDE-1 contains a conserved triad of acid aspartic residues (DDD) in its nucleotidyl transferase domain. Mutation of the corresponding DDD triad to DAD (D1011A) in human TUT4 resulted in loss of catalytic activity ^19^. A *cde-1* DAD mutant strain (Fig. 2a) showed similar viral susceptibility as the *cde-1* null mutants (Fig. 2b). In summary, we identify CDE-1-mediated 3′ terminal uridylation as an antiviral activity in the intestine of *C. elegans.*

**Figure 2.**
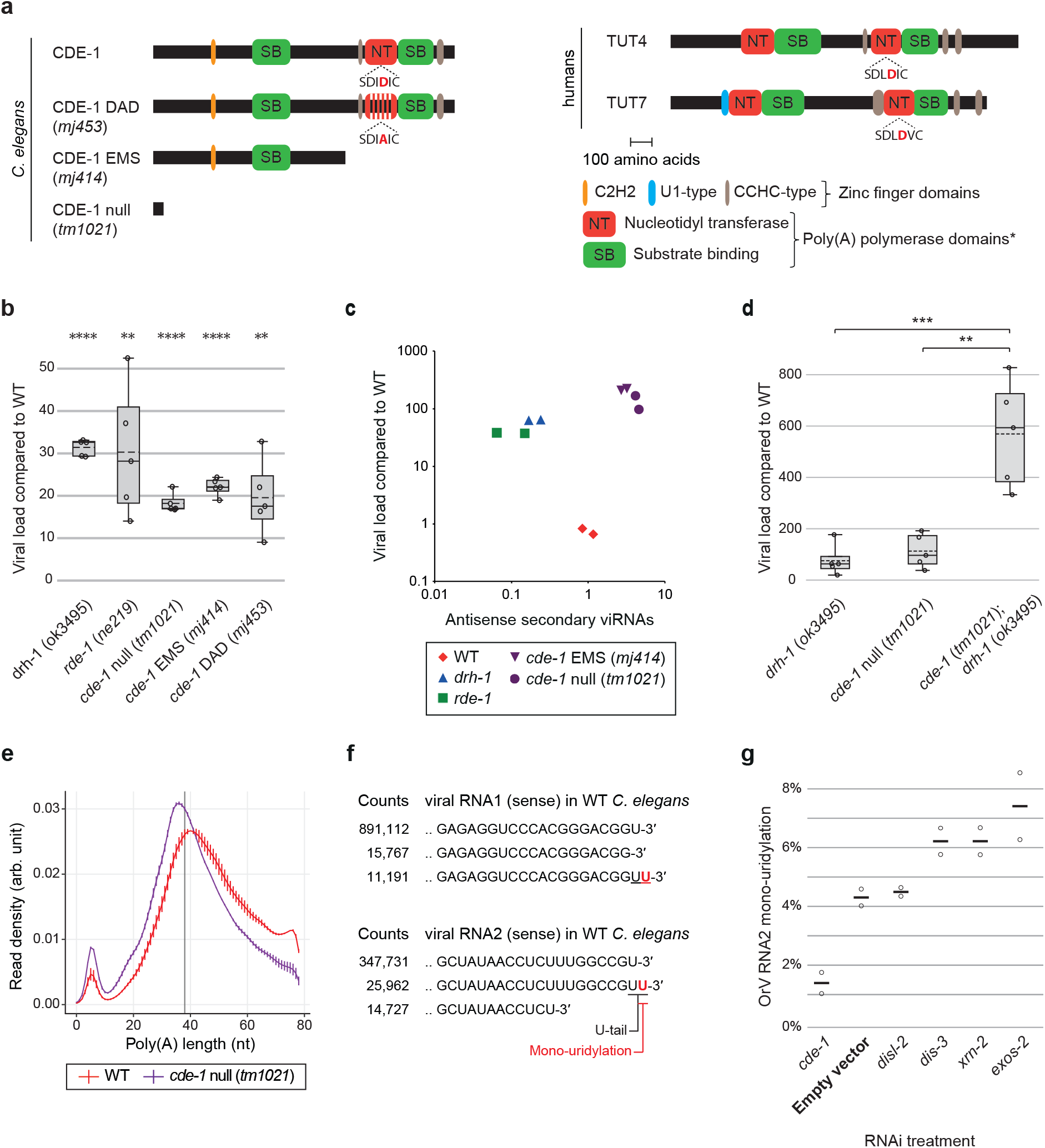
The terminal uridylyltransferase CDE-1 targets the Orsay virus RNA genome for uridylation. **a**, Diagrams of *C. elegans* CDE-1 and human TUT4 and TUT7. Domains were predicted by Interpro. The central D of the conserved DDD catalytic triad is highlighted in red. **b**, Viral load as measured by qRT-PCR of OrV RNA1 genome in adults two days after infection. Error: SEM, five biological replicates. One-tailed student’s t-test: **** p<0.0001, *** p<0.001, **p<0.01 **c**, Comparison between the viral load and secondary viRNA populations. Secondary viRNAs (22-nucleotide long, starting with a G, from 5′ tri/monophosphate RNA sequencing). Each data point represents one biological replicate (population of animals). **d**, As in **b**. **e**, Poly(A) tail length distribution measured by TAIL-seq after two days of OrV infection. Vertical grey line represents the mean of *cde-1* and wild type peaks (38 nt). **f**, Most frequent collapsed reads after RACE-seq on OrV RNA1 and RNA2 (2 dpi), respectively. Non-templated residues (absent from the reference genome) are indicated in red. **g**, Percentage reads with a non-templated mono-uridyl residue at the OrV RNA2 3′ end, upon RNAi-mediated gene knockdown as indicated, 1dpi, two biological replicates.

In eukaryotes, addition of 3′ U-tails by TUTs is a degradation signal that can engage: (i) the XRN-family of exoribonucleases for 5′ to 3′ RNA decay; (ii) the 3′ to 5′ exoribonuclease DIS3L2; (iii) the 3′ to 5′ exosome complex ^20-23^. We sought to identify the RNA(s) targeted by CDE-1 in its antiviral role. CDE-1 is implicated in endogenous RNAi pathways that are restricted to the germline ^16^. Small RNA sequencing on whole animals revealed that siRNAs are targeted by CDE-1 for 3′ uridylation, miRNAs are occasionally targeted, and piRNAs are not targeted ^16^ (Extended Data Fig. 6a). The role of CDE-1 in small RNA function remains unclear as depletion of CDE-1 leads to only subtle changes in siRNA and miRNA steady state levels (Extended Data Fig. S6b, c). To understand if CDE-1 functions through modification of siRNAs in antiviral immunity, we tested *cde-1* mutants directly for defects in antiviral RNAi. During an antiviral RNAi response in *C. elegans*, the ViRC complex recognizes the dsRNA of the replicating viral genome and dices it into sense and antisense ~23-nt long primary viRNAs, which are loaded into the RDE-1 Argonaute protein ^5^ (Extended Data Fig. 1). The RNAi response is further amplified by RNA-dependent RNA polymerase (RdRP, RRF-1) generated 22-nt long antisense secondary viRNAs, with a 5′ triphosphate guanine (22G-RNAs), which are incorporated into secondary Argonaute proteins ^5^ (Extended Data Fig. 1). Thus, a high viral load should correlate with a high level of viRNAs. We measured primary and secondary viRNAs in different genetic backgrounds (Extended Data Fig. 6d and Fig. 2c, respectively). All the mutants tested *(drh-1, rde-1, cde-1)* accumulate high levels of the virus as compared to wild type. In *drh-1* mutants, primary and secondary viRNAs are depleted when compared to wild type, despite the increase in viral load. In *rde-1* mutants, primary viRNAs are abundant but secondary viRNAs are depleted, as in *drh-1.* In contrast, *cde-1* mutants accumulate both primary and secondary viRNAs to a level that correlates with the high viral load. To determine if viRNAs can silence viral amplification in *cde-1* mutants, we carried out epistasis analysis using null mutants of *drh-1, rde-1* and *cde-1* (Fig. 2d and Extended Data Fig. 6e). Both *cde-1;drh-1* and *cde-1;rde-1* double mutants showed an increase in viral load as compared to *drh-1* or *rde-1* on its own. We conclude that CDE-1 does not exert its immune function through the antiviral RNAi pathway.

In mammals, uridylation is coupled to poly(A) tail length where TUT4 and TUT7 preferentially uridylate mRNAs with short poly(A) tails (<25 nt) to facilitate their degradation ^24,25^. We thus assessed the impact of CDE-1 on endogenous mRNA poly(A) tail lengths and terminal nucleotide addition in infected wild-type or *cde-1* mutant animals using TAIL-seq ^24,26^. The *C. elegans* transcriptome revealed a bimodal distribution of poly(A) tail lengths, with a major peak of poly(A) tails of ~40 nt, and a second peak of poly(A) tails of ~10 nt (Fig. 2e; using our method we could not assess transcripts with poly(A) tails > 79 nt). In *cde-1* mutants, there is a shift of the major ~40 nt peak to ~36 nt and an increase in transcripts with shorter poly(A) (Fig. 2e). We infer that CDE-1 promotes the degradation of transcripts with short poly(A) tails in *C. elegans* too. However, CDE-1 had no global effect on the poly(A) tail distribution of OrV-induced stress response genes (Extended Data Fig. 7a). Also, the OrV-induced stress response was stronger in *cde-1* mutants than in wild-type upon infection (Extended Data Fig. 7b), reflecting the difference in viral load between these two strains. This indicates that CDE-1 is not required for the OrV-induced stress response. Although we cannot formally rule out that CDE-1 may regulate an endogenous target(s), the evidence indicates this is not CDE-1’s principal function in antiviral immunity.

Instead, we postulated that the viral RNA genome itself may be uridylated by CDE-1. To detect uridylated Orsay RNA degradation intermediates, we carried out 3′ rapid amplification of cDNA ends (RACE) followed by high-throughput sequencing of the OrV RNAs extracted from *C. elegans* two days postinfection (RACE-seq; Extended Data Fig. 8a, 9a). Mono(U) tails constituted the most abundant fraction of non-templated nucleotides detected at the 3′ end of both OrV RNA1 and OrV RNA2 (Fig. 2f and Extended Data Figs 8b, 9b). For both RNA1 and 2, U-tailing was lost in two independent *cde-1* mutant alleles. In contrast, *drh-1* and *rde-1* mutants showed similar levels of viral RNA U-tails to wild-type, indicating that U-tailing is independent of viral load and that CDE-1 is not in limited quantities (Extended Data Fig. 8c, 9c). OrV RNA1 and RNA2 have a terminal uridylyl residue in their genome such that the addition of an extra non-templated uridine by CDE-1 forms a UU termination (Fig. 2f), which is a signal for uridylation-dependent RNA decay ^20,22^. The two XRN paralogs in *C. elegans* (XRN-1 and XRN-2) and the exosome components (e.g. DIS-3, EXOS-2) are essential ^27,28^, and these RNA degradation pathways normally act redundantly on uridylated RNAs ^25^. We therefore subjected *C. elegans* to a short (24 hours) RNAi treatment to effect a partial knockdown of *cde-1,* the exonuclease *disl-2* (the *C. elegans* DIS3L2 homologue), the exosome components *exos-2* and *dis-3,* and the exonuclease *xrn-2.* Treated animals, which appeared superficially wild type, were infected with OrV for 24 hours. The frequency of U-tails in OrV RNA2 was measured by RACE-seq (Fig. 2g). ~4% of OrV RNA2 were uridylated in animals exposed to the empty vector control RNAi, compared to ~1% in *cde-1* knockdown. RNAi treatments against *disl-2* did not affect the U-tail frequency. We measured a 1.4 to 1.7 fold increase in U-tail frequency upon RNAi treatment against *exos-2, dis-3* and *xrn-2,* suggesting that these factors each contribute to the degradation of uridylated viral RNAs, in accordance with a study that shows that DIS3 and the exosome can degrade viral RNAs in *Drosophila* and human cells ^29^. We conclude that *C. elegans* uses uridylation of the OrV as an innate immune defense. This mechanism acts in parallel to antiviral RNAi to combat viral infection (Extended Data Fig. 1).

The U-tail modification is conserved in eukaryotes and could impact a broad range of viruses in a variety of hosts ^30^. We tested if U-tailing affects the replication of Influenza A virus (IAV), which can infect human and murine cells. The IAV genome consists of eight antisense RNA segments (viral RNAs, vRNAs) from which the viral RdRP produces: (i) the sense complementary RNAs (cRNAs), which serve as templates to produce more vRNAs; and (ii) the mRNAs that are 3′ polyadenylated and exported to the cytosol for translation into viral proteins ^31^ (Extended Data Fig. 10a). We examined the 3′ end of a set of IAV RNAs, at 8 hours postinfection (hpi), in A549 human lung cells by RACE-seq. We could not detect U-tails at the ends of vRNAs or cRNAs (Fig. 3a and Extended Data Fig. 10b, c). In contrast, viral mRNAs were highly uridylated at their 3′ end, with ~77% of the IAV Nucleoprotein (NP) mRNA containing a U-tail, and a di(U)-tail being the most common type of 3′ end (~32%) (Fig. 3a and Extended Data Fig. 10d). The IAV NP mRNA is also uridylated (~40-50%) at 8 hpi in mouse embryonic fibroblasts (MEFs), but uridylation was lost in MEFs deficient in both *Tut4* and *Tut7* ^24^ (Fig. 3b). Thus TUT4/7 can uridylate the 3′ end of viral RNAs in mammals. The RACE-seq can only detect IAV mRNAs with poly(A) tails of <70 nt; it is possible that some IAV mRNAs with very long poly(A) tails are less prone to be uridylated. To test the impact of TUT4/7 on IAV, we measured the quantity of NP mRNA by qRT-PCR in infected MEFs (Fig. 3c). The IAV NP mRNA accumulated more rapidly and to a higher level at the peak in *Tut4/7* KO cells (peak at 8 hpi) compared to WT cells (peak at 16 hpi) before decreasing later in infection (24 hpi). Consistent with the difference in mRNA levels, the NP mRNA-encoded viral nucleoprotein (NP) accumulated more rapidly in *Tut4/7* KO cells compared to WT during the first eight hours of infection (Extended Data Fig. 10e). Accordingly, more infected cells overall were observed in *Tut4/7* KO compared to WT (Fig. 3d). To summarize, TUT4/7 reduces the expression levels of IAV mRNAs during the early stages of IAV infection, leading to a decrease in viral protein levels and rates of infection.

**Figure 3.**
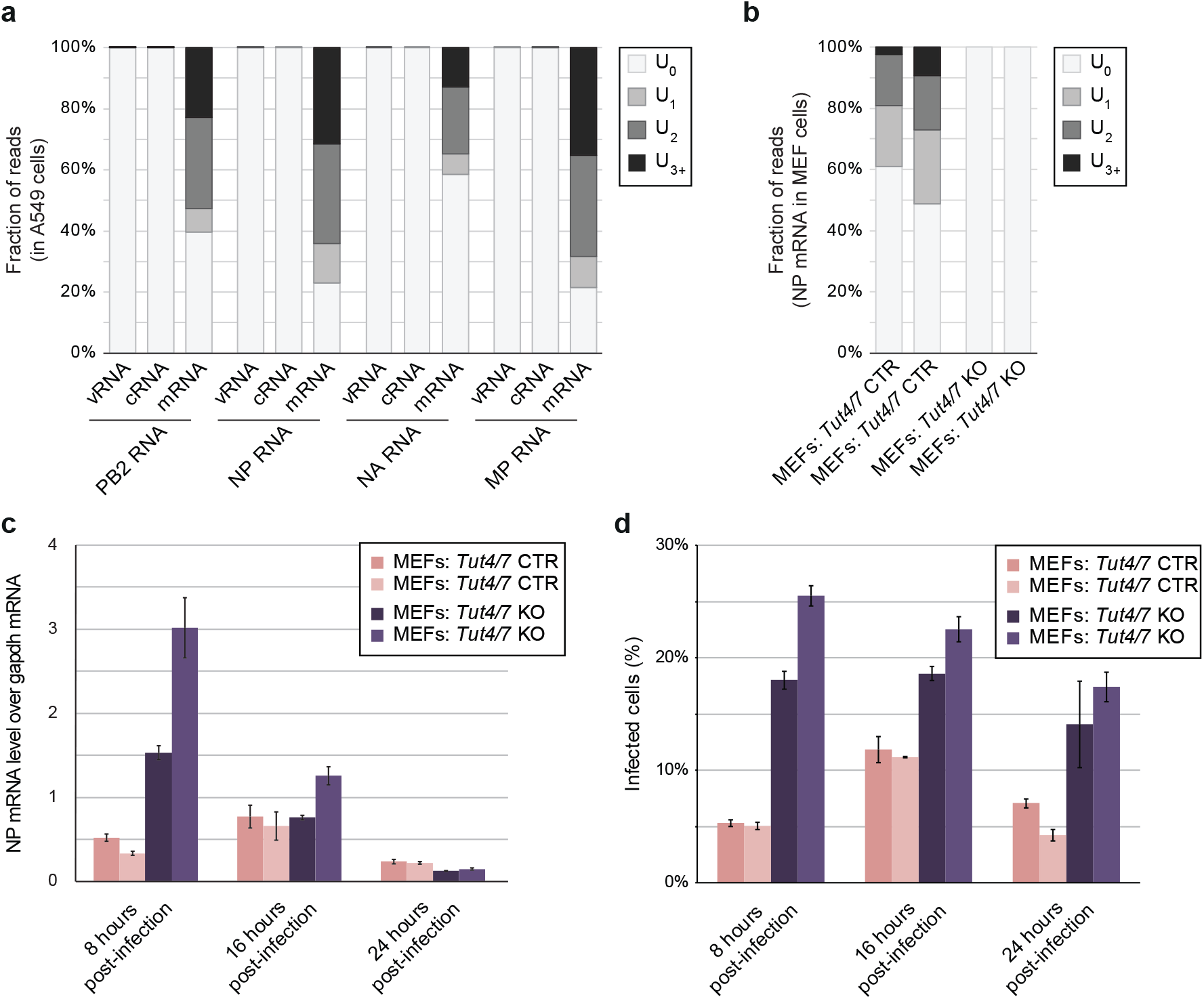
The terminal uridylyltransferases TUT4/7 attenuate Influenza A infection in mammals. **a**, Percentage of reads with a non-templated U-tail (no U-tail; 1 U; 2 Us or ≥ 3 Us) in different RNAs as indicated measured by RACE-seq in A549 cells 8 hpi. **b**, Percentage of reads with a non-templated U-tail (as in **a**) in MEF cells of different genotypes as indicated (with two independently created cell lines per genotype). **c**, Level of expression of the IAV NP mRNA normalized to *Gapdh* in MEF cells of different genotypes as indicated. Error: SEM, three biological replicates. **d**, Percentage of infected cells measured by immunofluorescence against NP (FACS). Error: SEM, three biological replicates.

Previously, we have shown that the antiviral RNAi pathway and DRH-1 are central to the innate immune response of *C. elegans* ^5^. Here, we demonstrate that the terminal uridylyltransferases also play a critical role in antiviral immunity, uridylating viral RNAs (with 1-2 Us) to mark them for degradation (Extended Data Fig. 1). It is unclear how terminal uridylyltransferase recognize viral RNAs as *bona fide* targets. Receptors of the RIG-I family commonly recognize pathogen-associated patterns at the 5′ termini of viral RNAs. In contrast, terminal uridylyltransferases interact with the 3′ termini of cytosolic RNAs with no poly(A)-tail or a short poly(A)-tail. As many RNA viruses, like OrV, lack a poly(A) tail at the 3′ termini of their RNA genomes, this may be a pathogen-associated pattern-recognition feature. We speculate that the IAV mRNAs and a fraction of the OrV RNAs are vulnerable to TUTs when exposed in the cytosol for translation. In conclusion, we find that terminal uridylyltransferases are potent antiviral factors during the early stages of RNA virus infections in *C. elegans* and in mammalian cells. This finding supports a scenario where eukaryotic mRNA decay pathways originally evolved as intrinsic cellular defenses against pathogens ^32,33^. Terminal uridylyltransferases are widely conserved in eukaryotes and could potentially target a wide range of RNA viruses ^30^. Our study illustrates that the 3′ termini of viral RNAs are key in the evolutionary arms race between viruses and their hosts.

## METHODS

### Genetics

Animals were grown on agar plates, at 20°C, and fed with *E. coli* strain HB101 (obtained from the *Caenorhabditis* Genetics Center, University of Minnesota, USA). Standard *C. elegans* procedures were used for maintenance and genetic crosses ^34^. The wild-type strain refers to Bristol N2 unless stated otherwise. All strains used in this study are listed in the Table S2.

### PCR primers

All PCR primers used in this study are listed in the Table S3.

### Viral filtrate preparation

Viral filtrate was prepared as in ^8^. Briefly, JU1580 animals were first stably infected by the Orsay virus (OrV) in solid culture and then transferred in a liquid culture containing OP50 bacteria for seven days. The liquid culture with infected JU1580 was then centrifuged at 16,000 g for 30 min and the supernatant was filtered (0.22 μm filter) to produce the viral filtrate (stored at -80°C).

### Transgenesis of *C. elegans* with the *lys-3p::GFP* viral stress sensor

The 452 bp region upstream of the *lys-3* start codon and the first 57 bp of the coding region of *lys-3* were used as a promoter and cloned into an entry clone using Multi-Site Gateway cloning (Invitrogen) according to manufacturer’s instructions. The *lys-3* donor plasmid was validated by sequencing. Gateway technology was then used to clone the *lys-3* fragment in frame with a GFP cDNA. The 3′ UTR of the *tbb-2* (tubulin, beta) gene was used. The *lys-3p::GFP:tbb-2-3′UTR* plasmid was amplified and purified according to Invitrogen’s instruction. The *C. elegans* microinjection mix was: 5 ng/μl plasmid *lys-3p::GFP:tbb-2-3′UTR;* 5 ng/μl co-injection marker *(myo-2::mcherry::unc-54-3′UTR,* pharynx expression) and 85 ng/μl 1 kb Invitrogen ladder in 1× injection buffer (20 mM potassium phosphate, 3 mM potassium citrate, pH 7.5). This mix was microinjected into the gonads of *rde-1 (ne219)* mutants to generate a multicopy extrachromosomal array (allele *mjEx547).* X-ray integration of the transgene into the *C. elegans* genome was performed as described previously ^35^. Animals carrying an integrated transgene (allele *mjIs228)* were outcrossed three times to generate SX2635 (lacking *ne219),* referred to as wild-type viral stress sensor strain in this study.

### Confocal images of the biostress reporter

A 2% agar pad was used on top of a glass slide and a drop of 10 μM tetramisol in M9 medium was placed on this agar pad. Animals were picked into the tetramisol solution. Imaging was performed with an Olympus Upright FV1000 microscope at 10× or 20× magnification, as specified, using the FluoView image software (Olympus). Identical microscope settings were used for all images within a figure.

### Forward genetic screen for Ovid screen isolates

Approximately 4,000 viral stress sensor transgenic animals were mutagenized using ethyl methanesulfonate (EMS) as described in ^34^ and ^36^. Approximately 50,000 F2 animals were infected for 3-4 days and ~2,000 animals showing intestinal GFP were picked individually for re-testing. 16 F2 families showed transmission of the viral stress sensor activation. Bleach treatment confirmed that removing OrV lead to a loss of intestinal GFP signal.

### *C. elegans* infection by the Orsay virus

Animals were either infected for four days as asynchronous populations or for two days as synchronous populations. Infections of asynchronous populations were performed as in ^5^. Briefly, two L4 hermaphrodites were distributed in each 50 mm plates and, on the next day, 20 μl of viral filtrate was spread on the plates. Animals were harvested (for viral load measurement) or observed under a Leica M165 FC fluorescent microscope (for scoring of the viral stress sensor) four days post-infection (4 dpi). This method was typically used for the characterization of the Ovid screen isolates. For the infection of synchronous populations, 200 animals at the larval stage L1 were deposited on each 50 mm plate. On the next day, L2 animals were infected with 20 μl of viral filtrate homogeneously spread on the plate. Plates were kept up-side-up for 24 hrs. Animals were harvested for viral load measurement at 2 dpi. This method was used to measure the viral load in *cde-1* mutants, as indicated in the figure legends.

### RNA level measurement by qRT-PCR

Harvested animals were washed three times by pelleting-resuspension in M9 solution. Lysis and qRT-PCR was then performed from 5 μl of animal pellet using the Power SYBR Green Cells-to-Ct kit (Ambion, Austin, TX) as described in ^5^. The primers M1835 and M1836 ^13^, and M4410 and M4411 ^4^, were used to measure RNA levels of *gapdh* and OrV gRNA1, respectively.

### RNAi-mediated knockdown of *unc-22*

All the bacterial feeding clones used in this study were a kind gift from the laboratory of Julie Ahringer. Bacteria were grown in LB-Ampicillin (50 μg/ml) for 6 hrs, then seeded onto 50 mm NGM agar plates containing 1 mM IPTG and 25 μg/ml Carbenicillin at a volume of 300 μl bacterial culture per plate and left to dry at room temperature, protected from the light, for 48 hrs. Two L4 animals were picked onto each RNAi plates and the young adult progeny were scored for the phenotype of interest after five days.

### Transgenesis of *C. elegans* with the CDE-1::GFP fosmid and imaging

The modified fosmid WRM064A_D06 where the GFP sequence is added at the N-terminal end of *cde-1* was provided by the TransgeneOme Project (Max Planck Institute of Molecular Cell Biology and Genetics, TransgeneOme Unit, Pfotenhauerstr. 108, 01307 Dresden, Germany; construct 09318202437763223 H08) ^37^. The construct was injected into the gonad of N2 animals to produce an extrachromosomal array (as described for the biostress reporter), using a *myo-3p::mCherry::unc-54-3′UTR* construct as a co-injection reporter. Transgenic animals (strain SX3123; allele *mjEx594)* were imaged with an Olympus Upright FV1000 microscope at 10x magnification.

### Fluorescence in situ hybridization of the Orsay virus RNA2

Animals were harvested in 15 ml of nanopure water and washed three times by pelleting-resuspension in nanopure water. Animals were then transferred to 1.5 ml tubes with a glass pipette. 1 ml of fixative solution (4% formaldehyde in 1X PBS) was added and samples were incubating at room temperature, on a rotating wheel, for 45 min. Nematodes were then washed twice by pelleting-resuspension in 1 ml of 1x PBS. Pellet of animals was resuspended in 1 ml 70% ethanol and stored at 4°C. After removal of the ethanol, fixed nematodes were washed once in 1 ml of wash solution (10% formamide, 2X SSC). The animal pellet was resuspended in 100 μl of hybridization solution (10% dextran sulfate, 2X SSC, 10% formamide) with 1 μl 1:50 of the probe v1580-RNA2-TexRed (ACCATGCGAGCATTCTGAACGTCA), a kind gift of Marie-Anne Félix, and incubated overnight at 30°C protected from the light. The next day, animals were washed three times in wash solution by pelleting-resuspension. Eventually, animals were resuspended in 1 ml wash solution with DAPI and incubated at 30°C for 30 min. Samples were centrifuged and supernatant was discarded. The animal pellet was resuspended in 1 ml of 2X SSC solution and stored at 4°C protected from light. Animals were then placed on a glass slide, in a drop of Vectashield anti-fade solution (Vector). Imaging was performed on an Olympus Upright FV1000 at 40x magnification, using the FluoView image software (Olympus). Same settings of fluorescence were used for all images compared.

### Transgenesis of *C. elegans* with the *vha-6p::gfp* plasmid and viral load measurement

The 878 bp region upstream of the *vha-6* start codon was used as a promoter and cloned into an entry clone using Multi-Site Gateway cloning (Invitrogen) according to manufacturer’s instructions. The *vha-6p* donor plasmid was validated by sequencing. Gateway technology was then used to clone the *vha-6p* upstream of (i) the GFP cDNA, or (ii) the full length *cde-1* gene (from ATG to STOP with endogenous introns). The 3′ UTR of the *tbb-2* (tubulin, beta) gene was used. The *vha-6p::GFP::tbb-2-3′UTR* and *vha-6p::cde-1::tbb-2-3UTR* plasmids were amplified and purified according to Invitrogen’s instruction. The *C. elegans* microinjection mix was: 10 ng/μl plasmid *vha-6p::GFP:tbb-2-3′UTR;* 10 ng/μl plasmid *vha-6p::cde-1::tbb-2-3′UTR;* 5 ng/μl co-injection marker *(myo-2::mcherry::unc-54-3′UTR,* pharynx expression) and 75 ng/μl 1 kb Invitrogen ladder in 1× injection buffer (20 mM potassium phosphate, 3 mM potassium citrate, pH 7.5). This mix was microinjected into the gonads of *cde-1 (tm1021)* mutants to generate a multicopy extrachromosomal array (allele *mjEx595). vha-6p* driven GFP expression was only observed in the intestine. 100 animals carrying the extrachromosomal array were manually selected for infection (from the L2 larval stage to young adult).

### Small RNA sequencing

Small RNA libraries were prepared from infected animals as previously described in ^5^. We used pellets of animals, washed three times in M9 solution and resuspended in 1 ml of TriSure (Bioline) as a starting material. RNA extraction was performed according to manufacturer’s instructions. Some populations of siRNAs (including secondary viRNAs) contain a characteristic 5′ triphosphate group that has to be replaced by a 5′ monophosphate to allow the 5′ ligation step of the library preparation. For this purpose, 1 μg of RNA was put in solution with 1X 5′p polyphophatase buffer and 1 μl of 5′ polyphophatase (Epicentre) for a total volume of 20 μl, incubated for 30 min at 37°C and then submitted to phenol purification and resuspended in 5 μl of nuclease-free water. Treated RNA sample was entirely used as starting material for the TruSeq Small RNA kit (Illumina), following the manufacturer’s instructions, to make the so-called 5′ independent libraries. So-called 5′ dependent libraries were made by a similar procedure but without polyphophatase treatment, so that only 5′ monophosphate siRNAs (such as primary viRNAs) could be cloned. Libraries were submitted to the Gurdon Institute sequencing facility for Illumina HiSeq sequencing (SR36). Small RNA sequencing data was aligned to the Ensemble WBcel235 release of the *C. elegans* genome using STAR ^38^ (v2.5.1b). Briefly, the aligner will allow untemplated residues at the ends of an aligned sequence when run in local mode. Untemplated 3′ sequences were extracted and analysed using custom Python scripts. Details of the analyses for each small RNA subtype can be found in the source code. For miRNA differential expression, reads were counted against the miRBase miRNA annotations (miRBase21 hairpins, WBcel235 genome) using featureCounts ^39^ (v1.5.0-p1). Differential expression analysis was performed on the counts using DESeq2 ^40^ (v1.10.1).

### CRISPR/Cas9 for *cde-1* catalytic dead mutant

A CRISPR/Cas9-mediated mutation of *cde-1* was generated as previously described ^41^. Guide RNA: UUUGCUGUCAAAUCCUUUGG. Homologous recombination template: TCAGCTATTGCTATTTGTTTGAGATTCGGAGATGGAGATGTTCCGCCTAAAGACTTGACAGCAAAAGAAGTTATTCAGAAAACTGAATCCGTTCTCAGAAAATGTCATTT. Only the D1069A missense mutation was introduced, as verified by sequencing.

### TAIL-seq

The TAIL-seq was performed as previously described in ^26^. Tail-seq libraries were processed using Tailseeker 2 ^26^. The 5′ and 3′ libraries were subsequently adapter trimmed using cutadapt 1.10 ^42^ with Illumina small RNA-seq adapters and filtered to a minimum length of 5bp. Trimmed 5′ reads were mapped with STAR 2.5.2a ^38^ against a combined meta-genome consisting of the *C. elegans* reference genome WBcel235 ^43^ and the OrV genome ^4^. Mapping was performed in end-to-end mode allowing no mismatches and a gap opening and extension penalty of 10,000. Reads were assigned to genes with bedtools 2.26.0 ^44^. Subsequently, 3′ reads without poly(A) tail or too many dark cycles were removed from the data. For the subsequent analysis, all *C. elegans* tags with poly(A) tail length equal to zero were discarded. Average poly(A) tail lengths and uridylation lengths for each sample were calculated as the arithmetic mean weighted by the support for each tag, reported by Tailseeker 2. The complete code is at <u>https://github.com/klmr/poly-u/tree/submitted</u>.

### mRNA libraries for deep sequencing

mRNA libraries were prepared from three biological replicates per sample, using the NEBNext Ultra RNA non-directional Library kit with poly(A) selection (NEB), according to manufacturer’s instructions. Libraries were submitted to the Gurdon Institute sequencing facility for Illumina HiSeq sequencing (SR30). Differentially expressed genes were then called using EdgeR ^45^.

### 3′ RACE-seq on the Orsay virus RNAs

The 3′ RACE was performed on the same RNA input than that used for small RNA libraries, without polyphosphatase treatment. 200 ng of RNA were submitted to 3′ ligation using the TruSeq Small RNA kit (Illumina), following the manufacturer’s instructions. 3′ ligated RNA was used for reverse-transcription, still using the TruSeq Small RNA kit whilst bypassing the 5′ ligation step. The 3′ end of OrV RNA1 (or RNA2) genome was amplified by PCR (“PCR1”) from 2 μl of cDNA, using the primers M7454 and M7456 (or M7455 and M7456) and the Phusion High-Fidelity Taq Polymerase (NEB) with CG buffer, according to manufacturer’s instructions. The thermocyler was programmed to 30 seconds at 98°C; 15 cycles of 5 seconds at 98 °C followed by 20 seconds at 60°C and 10 seconds at 72°C. The 5′ adapter sequence from the TruSeq Small RNA kit was then introduced at the 5′ end of the amplicons by PCR (“PCR2”) using the primers M7456 and M7601 for OrV RNA1 (or M7456 and M7602 for the OrV RNA2), using 2 μl of 1/10 diluted amplicon from PCR1 as a template and the same PCR conditions than that used in PCR1. The amplicons from PCR2 were purified using the DNA Clean & Concentrator-5 kit (Zymo Research) and resuspended in 10 μl of water. Resulting DNA was used as an input for the PCR amplification step of the TruSeq Small RNA kit, following the manufacturer’s instructions. Libraries were submitted to the Gurdon Institute sequencing facility for Illumina HiSeq sequencing (PE100). The libraries were run on a 10% polyacrylamide gel for size selection (the amplicons could be visualized under UV light and the bands were cut at the same distance of migration for all samples). Paired-end reads obtained from the 3′ RACE experiment on the viral genome show overlap. The PEAR software ^46^ was used to merge the paired reads into a single read (v0.9.6, default parameters). Merged reads not starting with the targeted 3′ viral genome sequence fragment were discarded. The targeted viral genome sequence was removed from the remaining reads using custom python scripts (<u>https://github.com/tdido/cde-1analysis</u>). The resulting sequences representing the untemplated tails were analyzed using custom python scripts.

### RNAi-mediated knockdown of *exonucleases*

Synchronized animals were grown on normal HB101 food until the L2 larval stage and then transferred RNAi food. Animals were left on RNAi plate (24 hours prior to infection) and infected for 24 hours, from the old L3/young L4 larval stages to adult. RACEseq was performed as described above.

### Cell culture

MEF cells were cultured with DMEM (GIBECO) supplemented with 12.5% FBS, 2mM L-glutamine, non-essential amino acid,100 units/ml penicillin/streptomycin, 100 uM β-mercaptoethanol (Sigma). Cells were splitted 1:4 and passaged every three days. A549 cells were cultured with DMEM (GIBECO) supplemented with 10% FBS, 2mM L-glutamine, non-essential amino acid,100 units/ml penicillin/streptomycin and 25mM HEPES.

### Generation of *Tut4/7* CTR and KO MEFs

*Tut4/7* CTR and KO MEFs were derived from E13.5 embryos from crosses of *Tut4^+/fl^;Tut7^+/fl^;R26^+/l+^* and *Tut4^+/fl^;Tut7^+/fl^;R26*^ERT-cre/ERT-cre^ mice by standard procedures and immortalized at passage 2 by two consecutive infections with pBabeSV40LT. Cre-mediated deletion to obtain Tut4/7 null alleles was induced with 600 nM 4-hydroxytamoxifen for three days ^24^.

### A549 and MEF cells infection by Influenza A virus and RACE-seq

Influenza A virus (A/WSN/1933, H1N1) used in this study was titrated on MDCK cells. All the inoculation MOI of influenza A virus described here and below was calculated as an equivalent MOI on the originally titrated MDCK cells.

A549 or MEF cells were trypsinized and seeded as 2X10^Λ^6 cells per T25 flask one day before infection. 16 hours after seeding, culture media were removed and cells were washed once with pre-warmed DMEM. Influenza A virus (A/WSN/1933, H1N1) were inoculated at MOI 3 diluted with 1000 μl DMEM supplemented with 0.1% BSA (D0.1B). Cells were trypsinized and collected 8 hours post infection. 750 μl TRIzol were added into each infected sample and were then freezed at -80 °C. RNA extraction was performed according to the standard TRIzol procedure.

For the A549 RACE-seq, 2 μg of RNA were submitted to 3′ ligation using the TruSeq Small RNA kit (Illumina), following the manufacturer’s instructions. 3′ ligated RNA was used for reverse-transcription, still using the TruSeq Small RNA kit (except that the Invitrogen Suprescript III was used instead of the Superscript II) whilst bypassing the 5′ ligation step. The RT final volume was 12.5 μl. After the RT, water was added to the samples to reach 18.5 μl, final volume. The 3′ end of IAV RNAs were amplified by PCR (“PCR1”) from 2 μl of cDNA, using the left primers M8443, M8444, M8451, M8452, M8453, M8454, M8455, M8456 (depending on the target, see the Table S3) with the right primer M7456 and the NEB Q5 polymerase, according to manufacturer’s instructions (25 μl reaction). The thermocyler was programmed to 30 seconds at 98°C; 5 cycles of 5 seconds at 98 °C followed by 20 seconds at 60°C and 20 seconds at 72°C. Each PCR product was purified using the DNA Clean & Concentrator-5 kit (Zymo Research) and eluted in 11 μl of water. The 5′ adapter sequence from the TruSeq Small RNA kit was then introduced at the 5′ end of the amplicons by PCR (“PCR2”) using the left primers M8459, M8460, M8467, M8468, M8469, M8470, M8471, M8472 (depending on the target, see the Table S3) with the right primer M7601, using 10 μl of purified PCR1 amplicon as a template and the same PCR conditions that used in PCR1. Again, the amplicons from PCR2 were purified using the Zymo columns and eluted in 11 μl of water. Resulting DNA was used as an input for the PCR amplification step of the TruSeq Small RNA kit, following the manufacturer’s instructions. Libraries were submitted to the Gurdon Institute sequencing facility for Illumina HiSeq sequencing (PE100). The libraries were run on a 10% polyacrylamide gel for size selection (the amplicons could be visualized under UV light and the bands were cut at the same distance of migration for all samples). Paired-end reads obtained from the 3′ RACE experiment on the viral genome show overlap. The PEAR software ^46^ was used to merge the paired reads into a single read (v0.9.6, default parameters). Merged reads not starting with the targeted 3′ viral RNA sequence fragment were discarded. The targeted viral genome sequence was removed from the remaining reads using custom python scripts (<u>https://github.com/tdido/cde-1_analysis</u>). The resulting sequences representing the untemplated tails were analyzed using custom python scripts. The MEFs RACE-seq was identical to the A549 cells RACE-seq, except: (i) the starting material was 1 μg, (ii) the Invitrogen Superscript II was used for the RT, (iii) PCR1 and PCR2 had 10 cycles each.

### MEFs infection by Influenza A virus and qRT-PCR

MEF cells were trypsinized and seeded as 8X10^Λ^4 cells per well of 24-well plate one day before infection. 16 hours after seeding, culture media were removed and cells were washed once with pre-warmed DMEM. Influenza A virus (A/WSN/1933, H1N1) were inoculated at MOI 3 diluted with 250 ul DMEM supplemented with 0.1% BSA (D0.1B). Cells were trypsinized and collected 8, 16 and 24 hours post infection. 350 ul TRIzol were added into each infected sample. RNA was extracted using Direct-zol™ RNA MiniPrep (Zymo Research) purification according to the manufacture’s protocol and was finally eluted into 60 ul RNase/DNase free water. The extracted RNA was subjected to strand specific qRT-PCR to quantify influenza virus replication as described in ^47^.

### MEFs infection by Influenza A virus and FACS assay

MEF cells were trypsinized and seeded as 1X10^4 cells per well of 96-well plate one day before infection. 16 hours after seeding, culture media were removed and cells were washed once with pre-warmed DMEM. Influenza A virus (A/WSN/1933, H1N1) were inoculated at MOI 3 diluted with 50 μl DMEM supplemented with 0.1% BSA (D0.1B). Inoculum was removed after 1 hour of incubation at 37 °C. The infected cells were cultured with MEF cell culture medium with 2.5% FBS. 8 hours post inoculation, culture media were removed and cells were trypsinized through incubation with 30 μl 0.05% trypsin for 3 minutes at 37 °C. Trypsinized cells were resuspended with 70ul of P2F (PBS with 2% FBS) and then fixed with 100 μl 4% PFA for 15 minutes. Fixed cells were centrifuged at 300g for 5 minutes and then washed once with 100 μl P2F. Cells were then permeablized with buffer (0.1% Saponin, 10mM HEPES, 0.025% Sodium Azide in 1XHBSS) for 15 minutes at room temperature and then spinned at 500g for 2 minutes to remove buffer. Primary anti-influenza A virus nucleoprotein antibodies were purchased from Millipore (MAB8258B | clone A3, biotin-conjugated). The primary antibodies were diluted 1:2000 in permeable buffer and 50 μl diluted antibodies were added into each well of 96-well plate. Primary antibodies were incubated with infected cells at room temperature for 1 hour. The cells were then washed 3 times with permeable buffer. FITC conjugated goat anti-mouse secondary antibodies were purchased from Invitrogen and diluted at 1:1000 in permeable buffer. Secondary antibodies were incubated for 1 hour at room temperature and washed as described before. The stained cells were finally resuspended in 70 μl P2F. The cell suspension was run on a high throughput FACS machine (MACSQuant^®^ analyzer 10 – Miltenyi Biotec). Uninfected cells were stained the same as infected cells and were used as negative staining cell populations. Any cells/events that had fluorescence intensity higher than all the negative staining cell population were gated as virus infection positive. Data were analyzed using flowjo software (version 10).

## ACCESSION CODES

All raw sequencing data are deposited in GEO (small RNA sequencing: GSE80169; mRNA sequencing: GSE76901; TAILseq: GSE85893). All *C. elegans* strains created in this study will be freely available on a non-collaborative basis. Correspondence and requests for materials should be addressed to E.A.M.

## ACKNOWLEDGMENTS

We thank Mélanie Tanguy for OrV viral filtrates, Lise Frézal for help with the OrV RNA FISH, Isabel Wilkinson for support with the genetic screen, Nicolas J. Lehrbach for help with microinjections, and Marc Ridyard for lab management. We thank Kay Harnish, Fabian Braukmann and Sylviane Moss for high-throughput sequencing support. We are grateful to V. Narry Kim and Hyeshik Chang for sharing information on TAIL-seq and Adrianus C.M. Boon for providing IAV. We thank the International *C. elegans* gene knockout consortium and the TransgeneOme project for providing reagents. We thank Vladimir Benes and the EMBL genome core for sequencing support. We thank George Allen and Charles Bradshaw for core bioinformatics support. This work was supported by Cancer Research UK (C13474/A18583, C6946/A14492), the Wellcome Trust (104640/Z/14/Z, 092096/Z/10/Z) and The European Research Council (ERC, grant 260688). DW holds an Investigator in the Pathogenesis of Infectious Disease Award from the Burroughs Wellcome Fund.

## AUTHOR INFORMATION

The authors have made the following declarations about their contributions: Conceived and designed the experiments: J.L.P., H.J., E.A.M. Performed the experiments: J.L.P., H.J., E.K., J.K., M.M., C.M.. Analyzed the data: J.L.P., H.J., T.D.D., K.L.M.R., A.J.E, D.O.C., D.W., EAM. Wrote the manuscript: J.L.P., E.A.M.

## EXTENDED DATA FIGURES

**Extended Data Figure 1.**
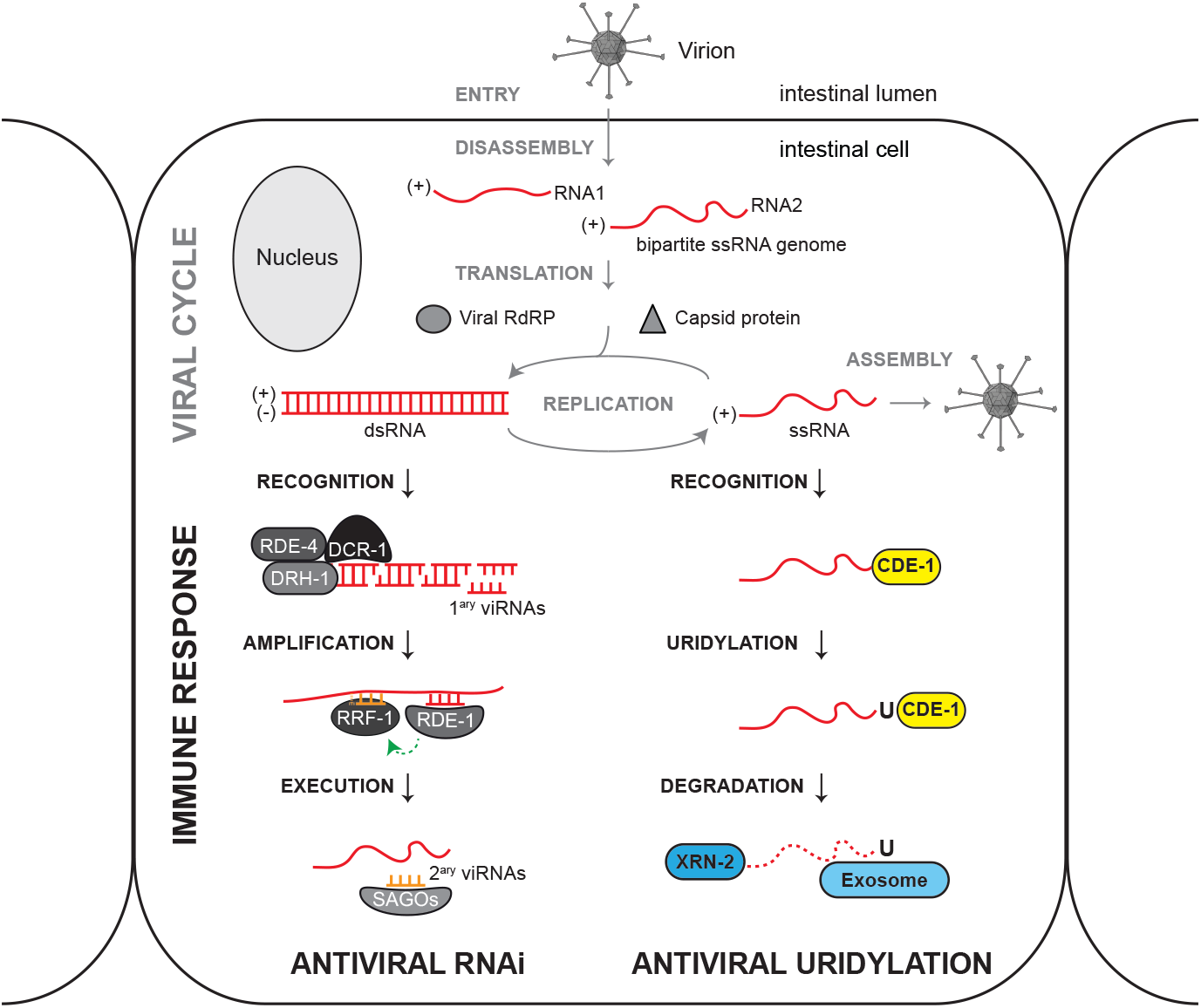
Antiviral RNAi and virus terminal uridylation are parallel immune defense pathways. Virion cartoon adapted from ^12^. The Orsay virus (OrV) is a bipartite positive-strand RNA virus related to the *Nodaviridae.* As is typical for positive sense RNA viruses, the genomic strand of OrV is a template for translation. OrV spreads horizontally in populations of *C. elegans:* it is taken up orally, infects only intestinal cells and probably exits through defecation. Antiviral RNAi pathway as in ^5^.

**Extended Data Figure 2.**
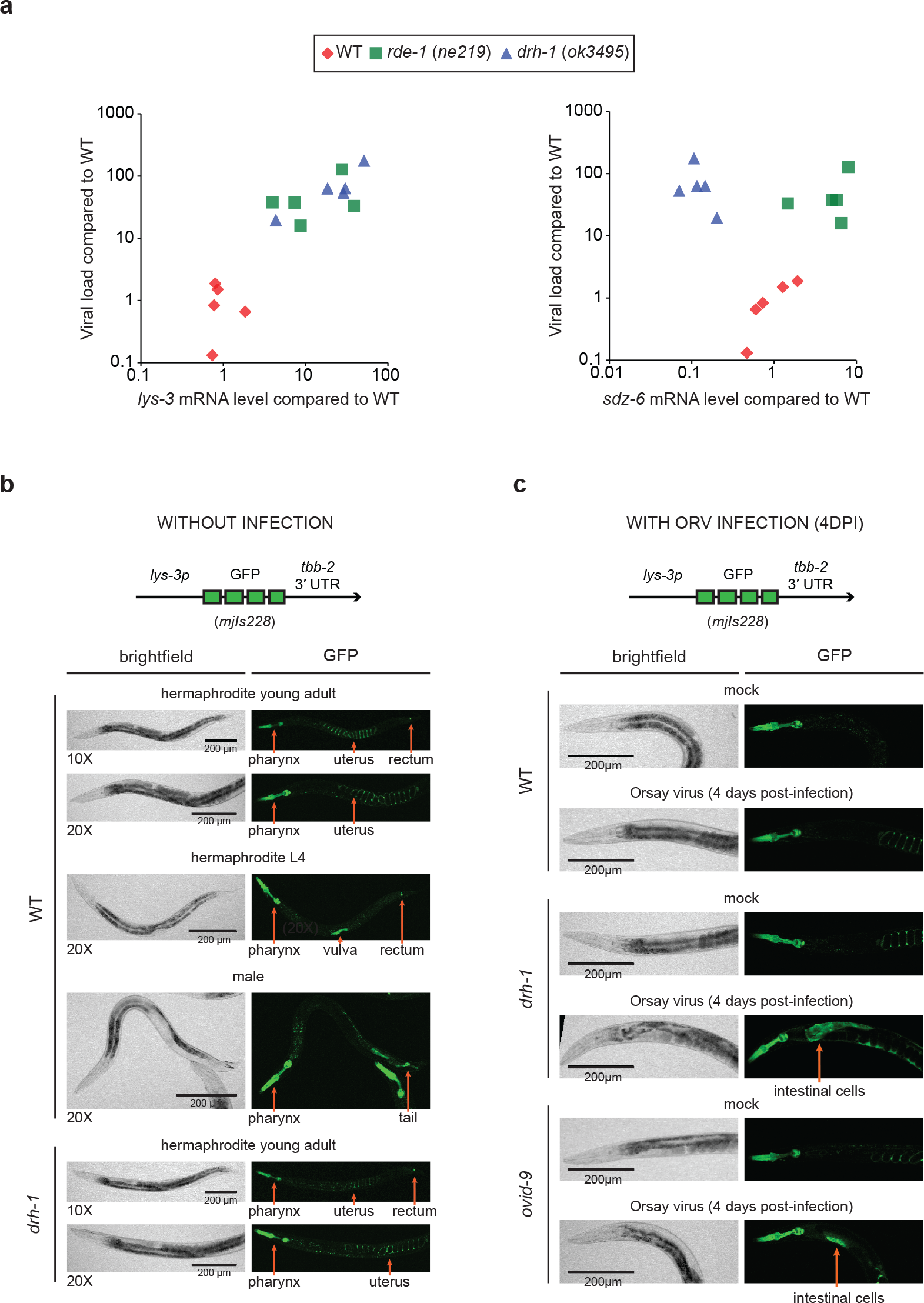
The viral stress sensor *(lys-3p::GFP)* is constitutively active in some tissues but is induced in the intestine upon severe viral infection. **a**, Comparison of viral load and the *lys-3* and *sdz-6* mRNA expression after two days of infection by qRT-PCR, strains as indicated. Each data point represents one biological replicate (population of animals on one agar plate). DRH-1 induces a virus-specific stress response upon OrV infection (e.g. *sdz-6).* However, the antiviral function of the DRH-1-mediated stress response remains to be elucidated. *C. elegans* also elicits a “biotic stress response” upon OrV infection that is independent of DRH-1 and partially overlaps with transcriptional responses induced by other types of pathogens, possibly as a result of perturbations in cell homeostasis and/or mechanical integrity (e.g. antibacterial lysozyme-3; *lys-3).* **b**, Representative confocal sections (10× or 20× magnification, as specified) of the viral stress sensor in wild type and *drh-1* mutants without infection. The viral stress sensor exhibited constitutive activity in uninfected individuals, which was restricted to specific tissues. GFP was observed at all developmental stages in the pharynx and the rectum of hermaphrodites. Additionally, hermaphrodites at the L4 larval stage would show a strong GFP signal around the vulva and gravid adults exhibited the GFP in the uterine lumen. In males, GFP was observed in the pharynx and the tail. GFP expression was comparable in wild type and *drh-1* mutants and independent of viral infection. Thus, the gene *lys-3* is constitutively active in tissues neighboring openings exposed to the environment, the most likely entry points of potential bacterial pathogens. **c**, Representative confocal sections (20× magnification) of young adults (strains as indicated) carrying the viral stress sensor. Animals were uninfected (mock) or infected with OrV for four days. The viral stress sensor was strongly induced in the intestine after infection of *drh-1* mutants, which is in agreement with the tropism of OrV. Intestinal GFP was most often visible around the collar of the nematodes, in the anterior region of the intestine in young adults. Some infected individuals exhibited a strong GFP signal throughout their entire body (data not shown), suggesting that the induction of the viral stress sensor can spread from cell to cell, like an inflammation process. The viral stress sensor offers an opportunity to easily monitor viral infections in living animals.

**Extended Data Figure 3.**
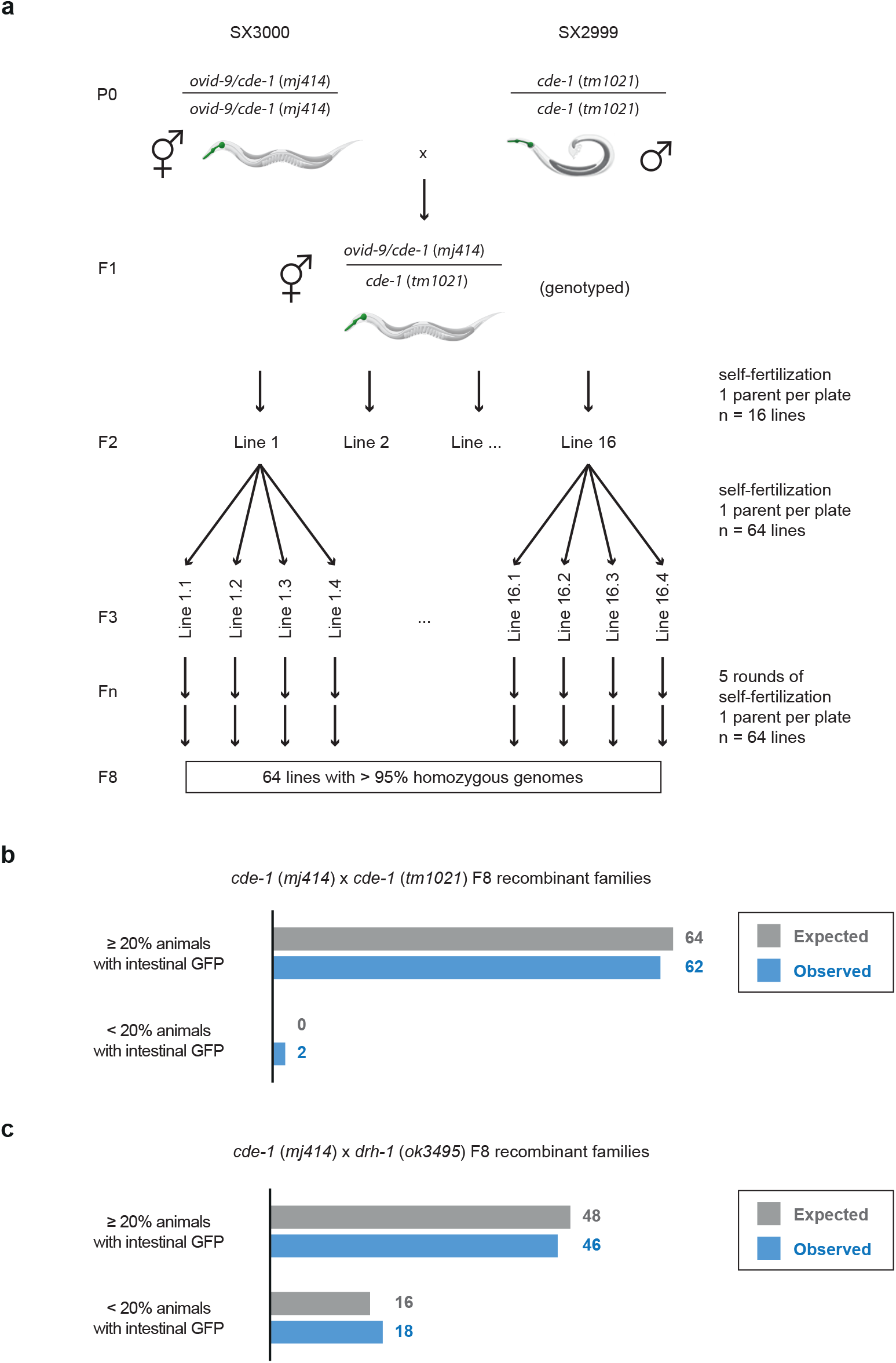
A *cde-1* deletion allele fails to complement the screen isolate *ovid-9*. **a**, Workflow of *cde-1/ovid-9 (mj414) × cde-1 (tm1021)* F8 recombinant family generation. A similar strategy was used to construct the *cde-1 (mj414) × drh-1 (ok3495)* F8 recombinant families. All animals were homozygous for the viral stress sensor *(mjIs228).* **b-c**, Number of families that activated the viral stress sensor in more than 20% of individuals after four days of infection with OrV. Approximately 50 individuals were scored per agar plate.

**Extended Data Figure 4.**
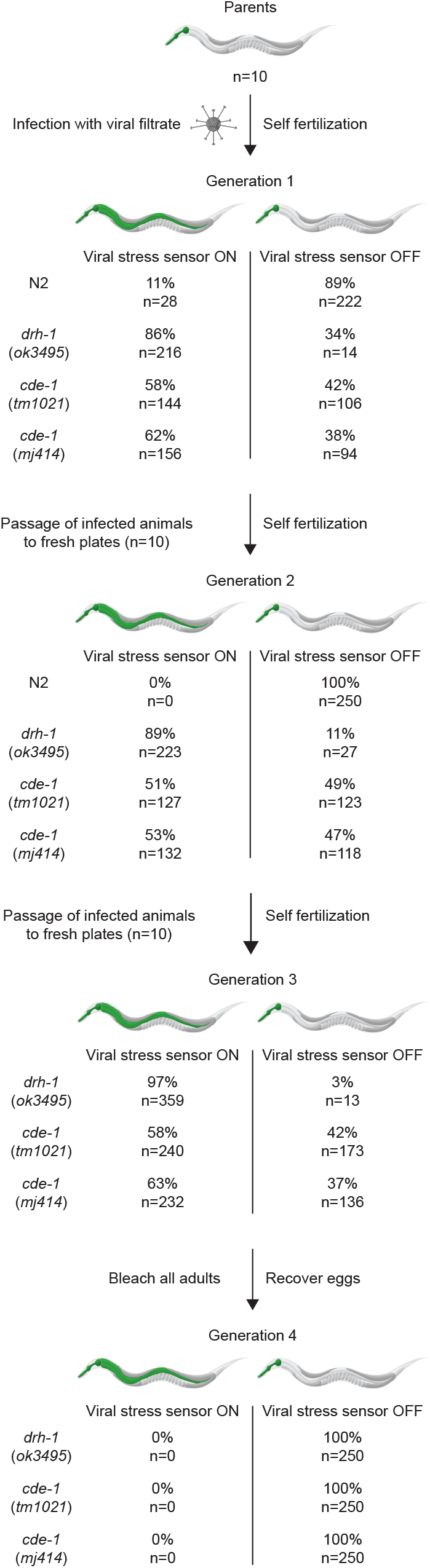
*cde-1* mutants show horizontal transmission of Orsay virus infection. Workflow and data monitoring the inter-individual transmission of OrV infection (in strains as indicated) using the viral stress sensor.

**Extended Data Figure 5.**
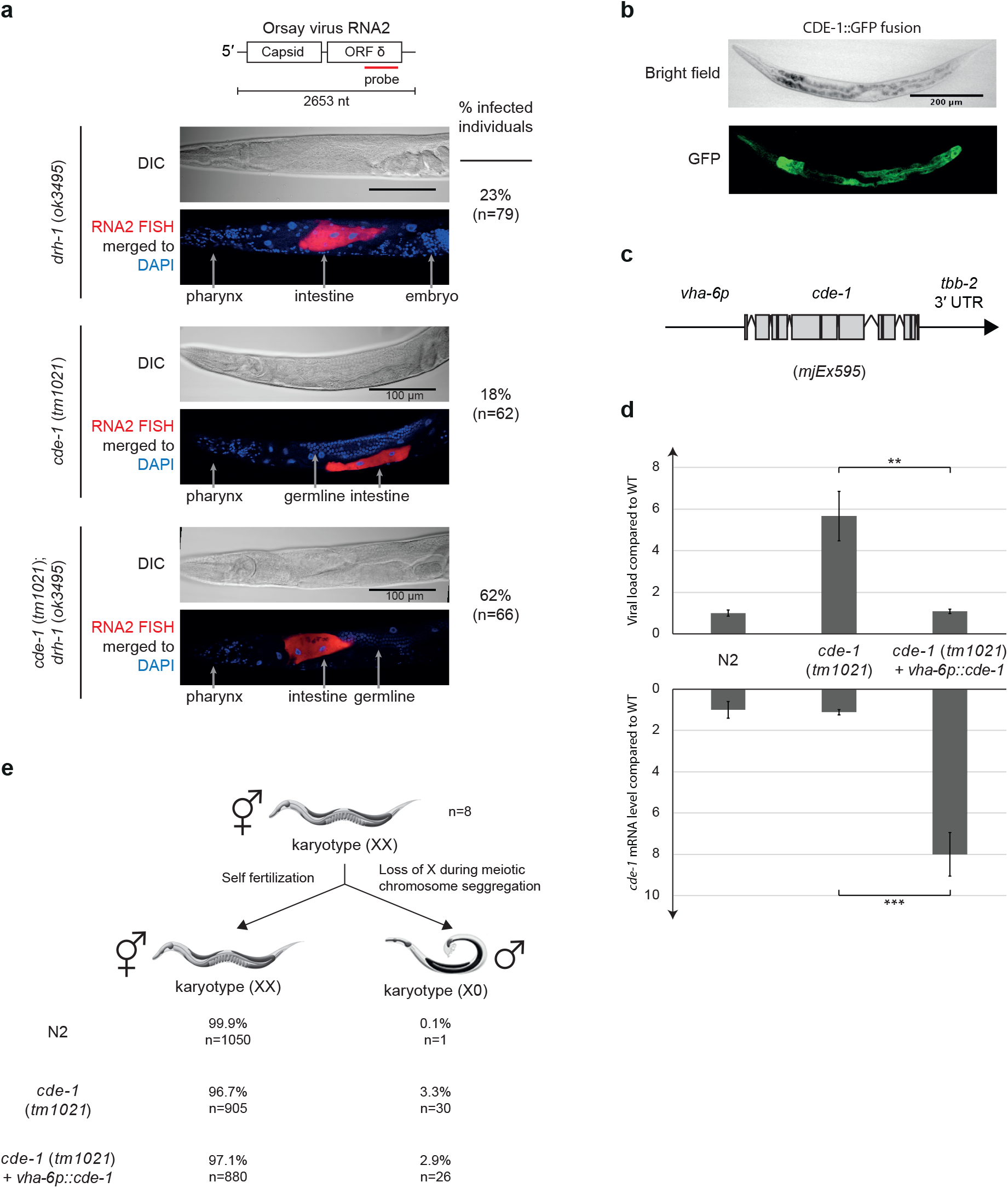
Intestinal expression of *cde-1* confers antiviral immunity. **a**, Representative confocal sections (20× magnification) of OrV *in vivo* RNA FISH. **b**, Representative confocal section (10× magnification) of a *C. elegans* L4 larva expressing *cde-1::GFP.* As two previous reports disagreed about the expression pattern of CDE-1 ^16,17^, we used fosmid-recombineering to generate transgenic animals driving GFP expression from an endogenous genomic context. **c**, Diagram of the *cde-1* rescue transgene, using the intestine-specific promoter of the gene *vha-6.* This transgene was injected in *cde-1* null mutants. **d**, Viral load as measured by qRT-PCR of OrV RNA1 genome in adults two days after infection. Error: SEM, five biological replicates. One-tailed student’s t-test: *** p<0.001, **p<0.01. **e**, Incidence of male in the progeny of hermaphrodites left to self-fertilize at 25°C, in strains as indicated.

**Extended Data Figure 6.**
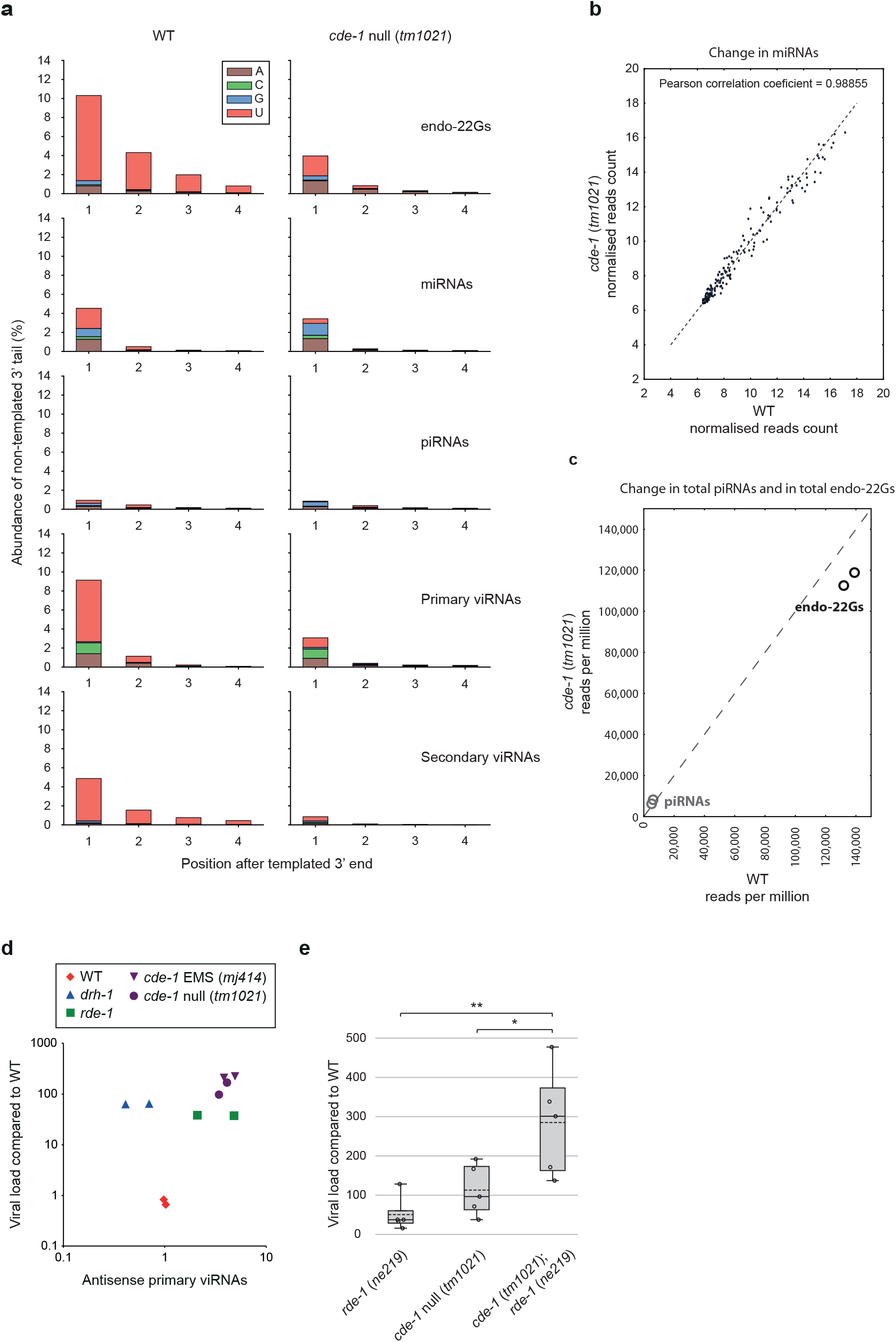
CDE-1 is not required for general miRNA homeostasis. **a**, Non-templated nucleotides at the 3′ end of the different classes of endogenous and antiviral small RNAs as indicated. RNA was isolated from young adults after two days of infection with OrV. **b**, miRNA expression in *cde-1 (tm1021)* mutants as compared to wild type, samples as in **a**. **c**, piRNAs and endogenous 22G-RNAs abundance in *cde-1 (tm1021)* mutants as compared to wild type, normalised to library size. Each data point represents one biological replicate. Samples as in **a**. **d**, Comparison between the viral load and primary viRNA populations. Primary viRNAs (23-nucleotide long, from 5′ monophosphate RNA sequencing). Only antisense RNAs were considered to exclude potential degradation products. Each data point represents one biological replicate (population of animals). Samples as in Fig. 2c. **e**, Viral load as measured by qRT-PCR of OrV RNA1 genome in adults two days after infection. Error: SEM, five biological replicates. One-tailed student’s t-test: **p<0.01, *p<0.05. Samples as in Fig. 2b.

**Extended Data Figure 7.**
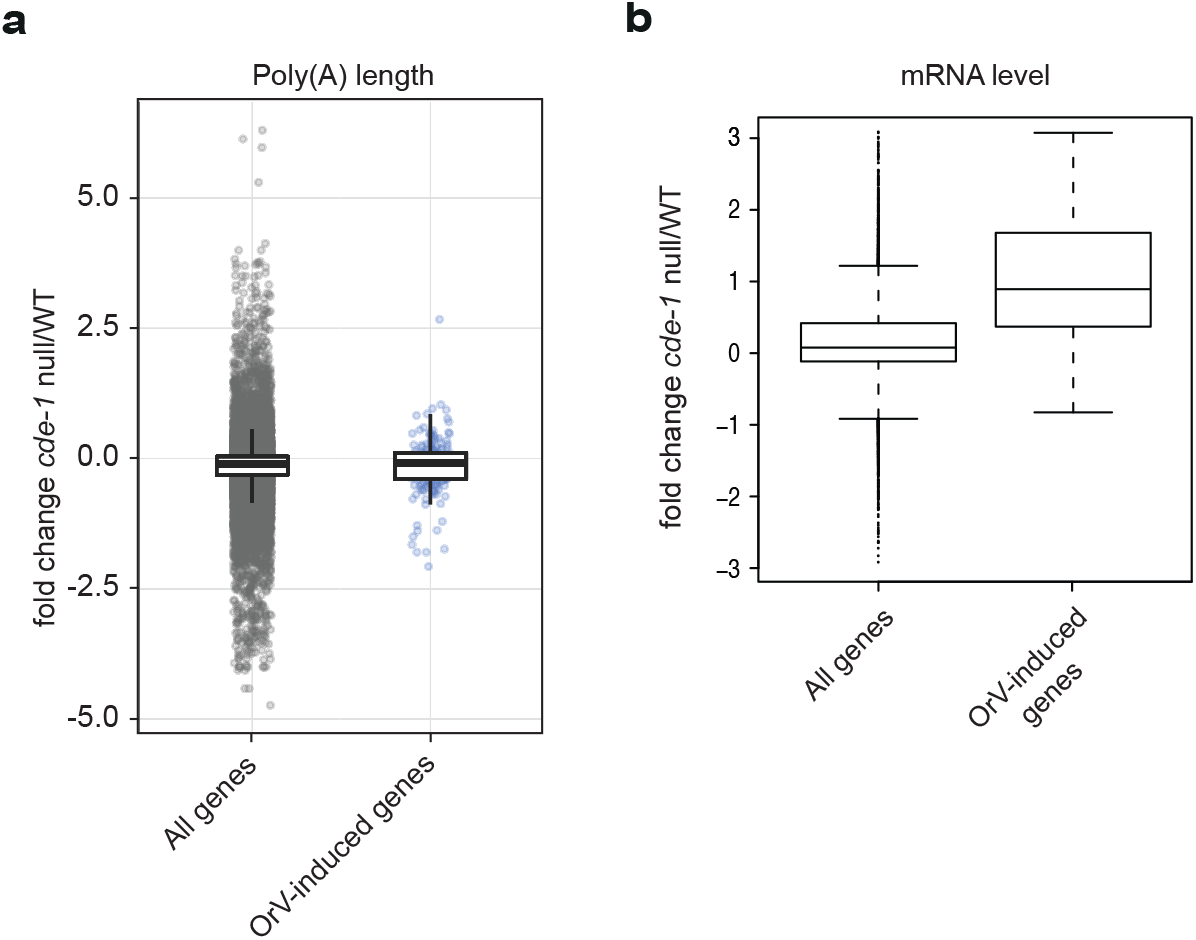
CDE-1-depleted animals show a high expression of stress response genes during Orsay virus infection. **a**, Fold change in the length of poly(A) tails (measured by TAIL-seq) in *cde-1* mutants compared to wild type. RNA was isolated from young adults after two days of OrV infection. Samples as in Fig. 3a. **b**, Differential mRNA expression in *cde-1 (tm1021)* compared to wild type, two days of OrV infection (mRNA-seq).

**Extended Data Figure 8.**
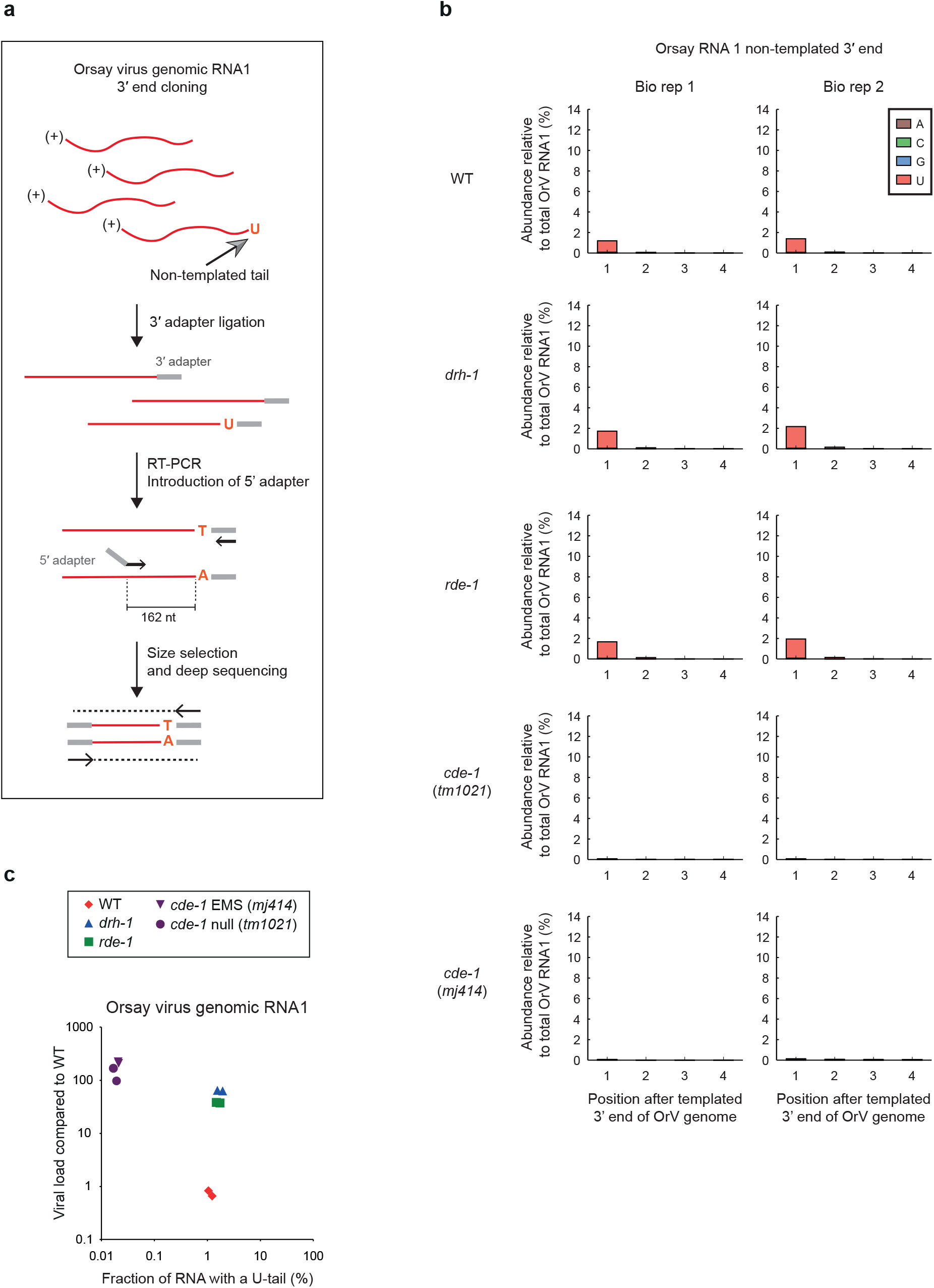
The 3′ end of the Orsay virus genomic RNA1 contains *cde-1* dependent non-templated U-tails in *C. elegans*. **a**, Simplified workflow of 3′ RACE-seq of OrV RNA1. **b**, Abundance of non-templated nucleotides detected at the 3′ end of OrV RNA1 in strains as indicated, two days of infection. Two biological replicates per genotype. **c**, Comparison between the viral load and the fraction of non-templated mono(U) tails at the 3′ end of OrV RNA1 in strains as indicated. Each data point represents one biological replicate. Samples as in Fig. 2c.

**Extended Data Figure 9.**
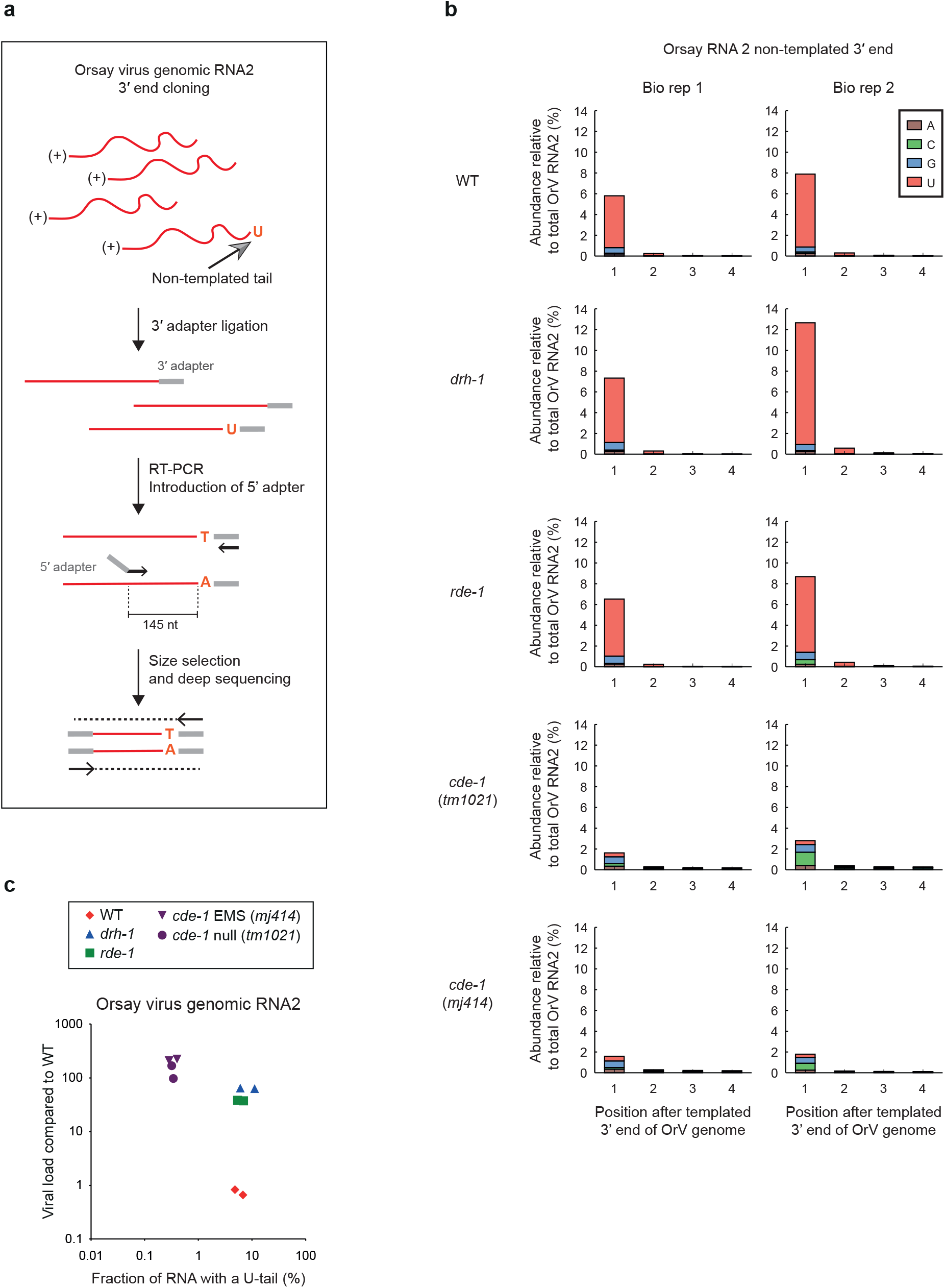
The 3′ end of the Orsay virus genomic RNA2 contains *cde-1* dependent non-templated U-tails in *C. elegans*. **a**, Simplified workflow of 3′ RACE-seq of OrV RNA2. **b**, Abundance of non-templated nucleotides detected at the 3′ end of OrV RNA2 in strains as indicated, two days of infection. Two biological replicates per genotype. Samples as in Extended Data Fig. 8b. **c**, Comparison between the viral load and the fraction of non-templated mono(U) tails at the 3′ end of OrV RNA2 in strains as indicated. Each data point represents one biological replicate. Samples as in Fig. 2c.

**Extended Data Figure 10.**
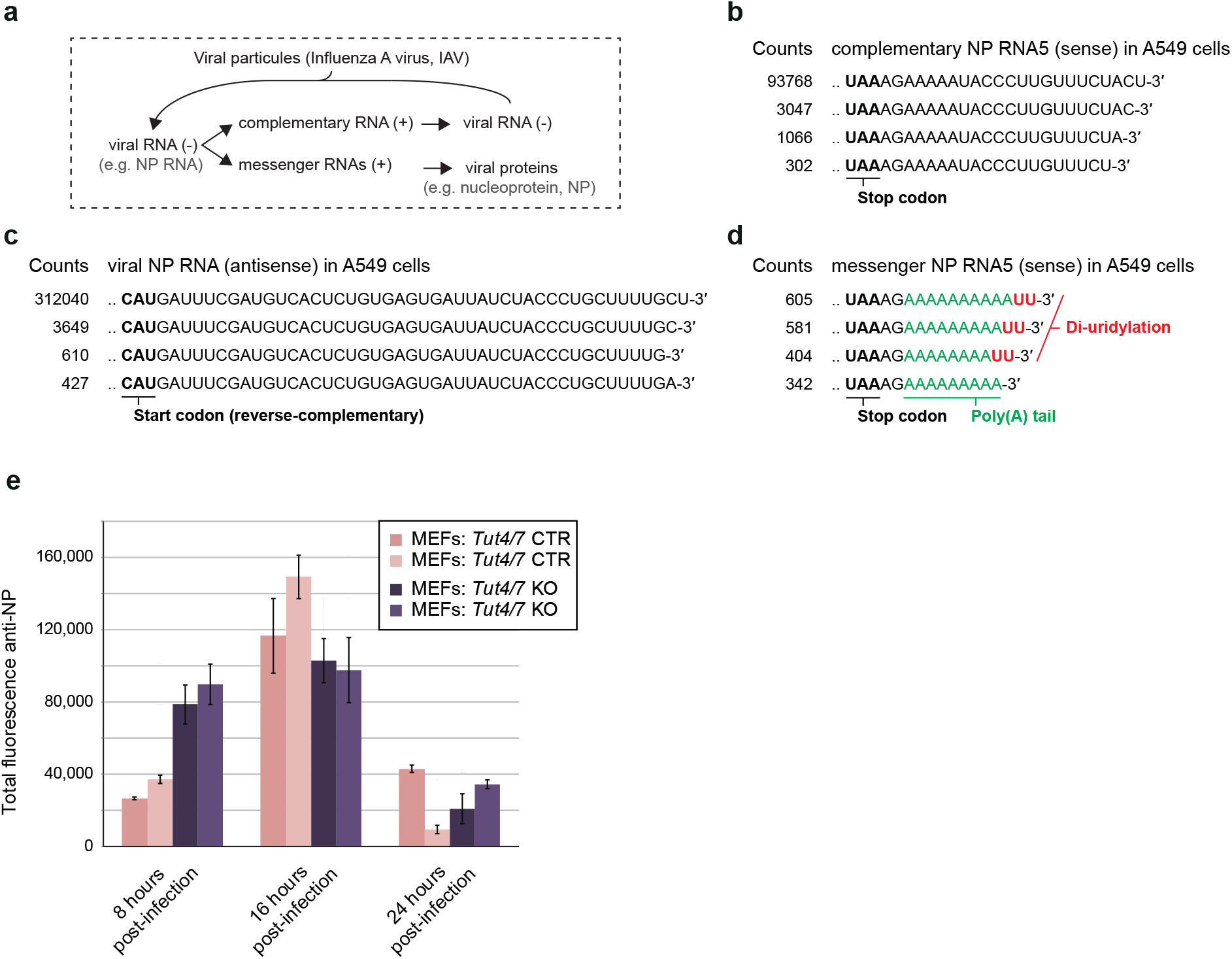
The terminal uridylyltransferases TUT4/7 target Influenza A mRNAs in mammals. **a**, Schematic of the Influenza A virus viral cycle. **b-d**, Most frequent collapsed reads after RACE-seq on IAV NP cRNA, NP vRNA and NP mRNA, respectively in A549 cells at 8 hpi. **e**, Protein level of the IAV NP measured by immunofluorescence (FACS). Error: SEM in three biological replicates.

## SUPPLEMENTARY INFORMATION

**Supplementary Table 1.** Infection by the Orsay virus induces viral stress sensor in Ovid screen isolates.

| Strain | Genotype | % intestinal<br>GFP mock | % intestinal<br>GFP OrV |
| --- | --- | --- | --- |
| SX2635 | WT | 0 | 4 |
| SX2615 | *rde-1* | 0 | 42 |
| SX2790 | *drh-1* | 0 | 83 |
| SX2996 | *ovid-1* | 0 | 86 |
| SX2900 | *ovid-2* | 0 | 83 |
| SX2739 | *ovid-3* | 0 | 74 |
| SX2901 | *ovid-4* | 0 | 73 |
| SX2902 | *ovid-5* | 0 | 68 |
| SX2729 | *ovid-6* | 0 | 64 |
| SX2991 | *ovid-7* | 0 | 60 |
| SX2990 | *ovid-8* | 0 | 52 |
| SX3000 | *ovid-9* | 0 | 48 |
| SX2881 | *ovid-10* | 0 | 47 |
| SX2987 | *ovid-11* | 0 | 46 |
| SX2988 | *ovid-12* | 0 | 44 |
| SX2920 | *ovid-13* | 0 | 26 |
| SX2909 | *ovid-14* | 0 | 26 |
| SX2886 | *ovid-15* | 0 | 24 |
| SX2885 | *ovid-16* | 0 | 24 |

**Supplementary Table 2.**
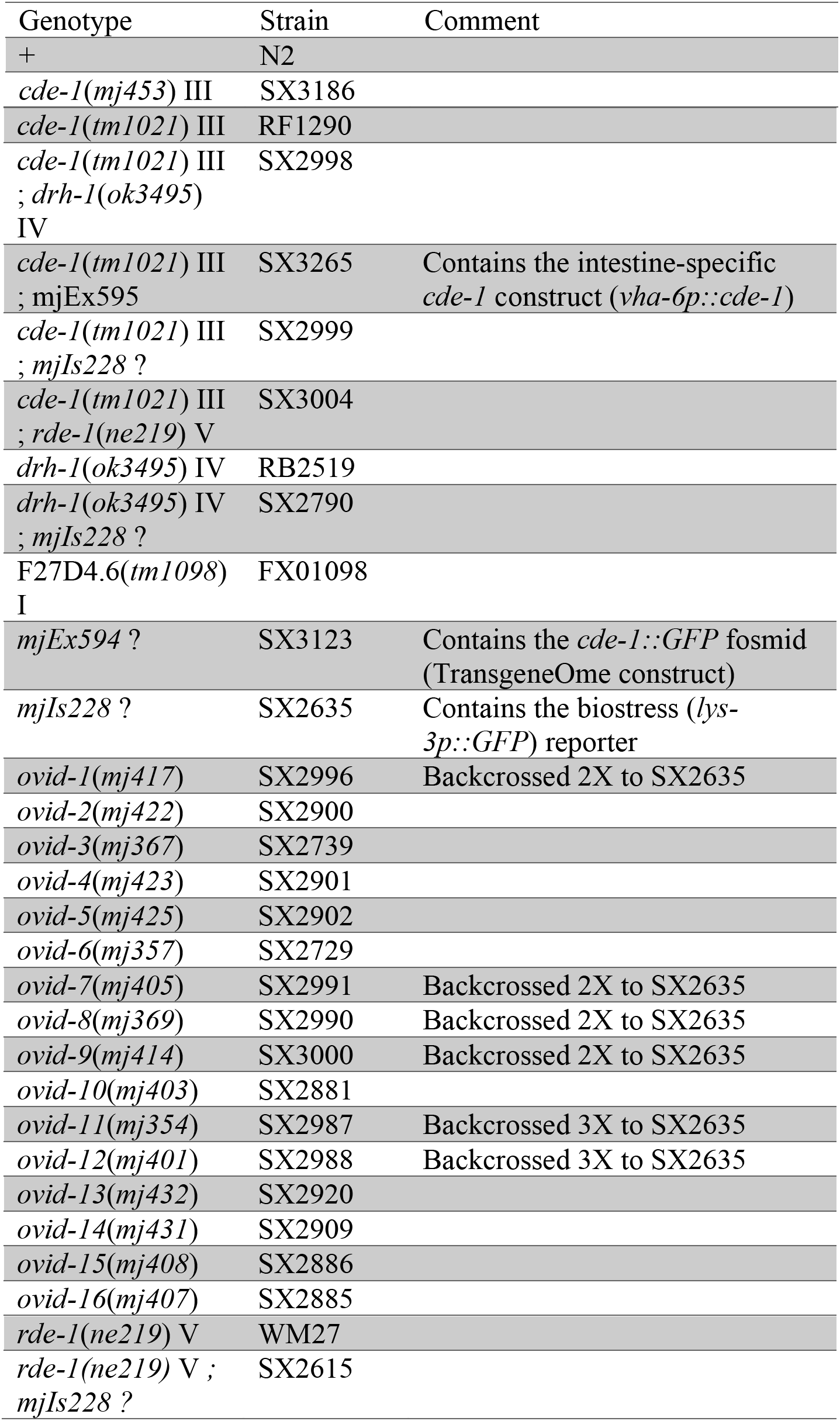
*C. elegans* strains used in this study

**Supplementary Table 3.**
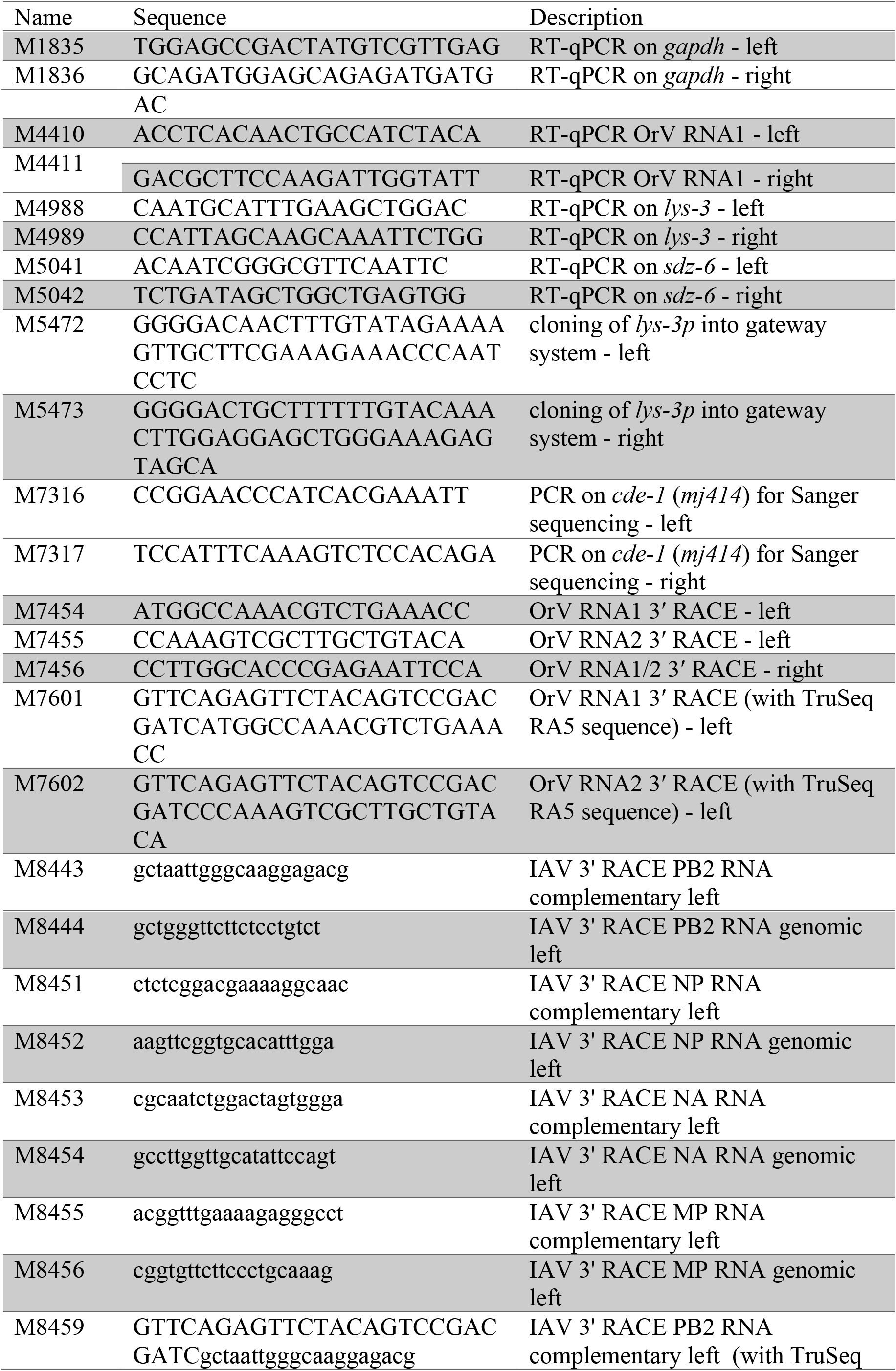
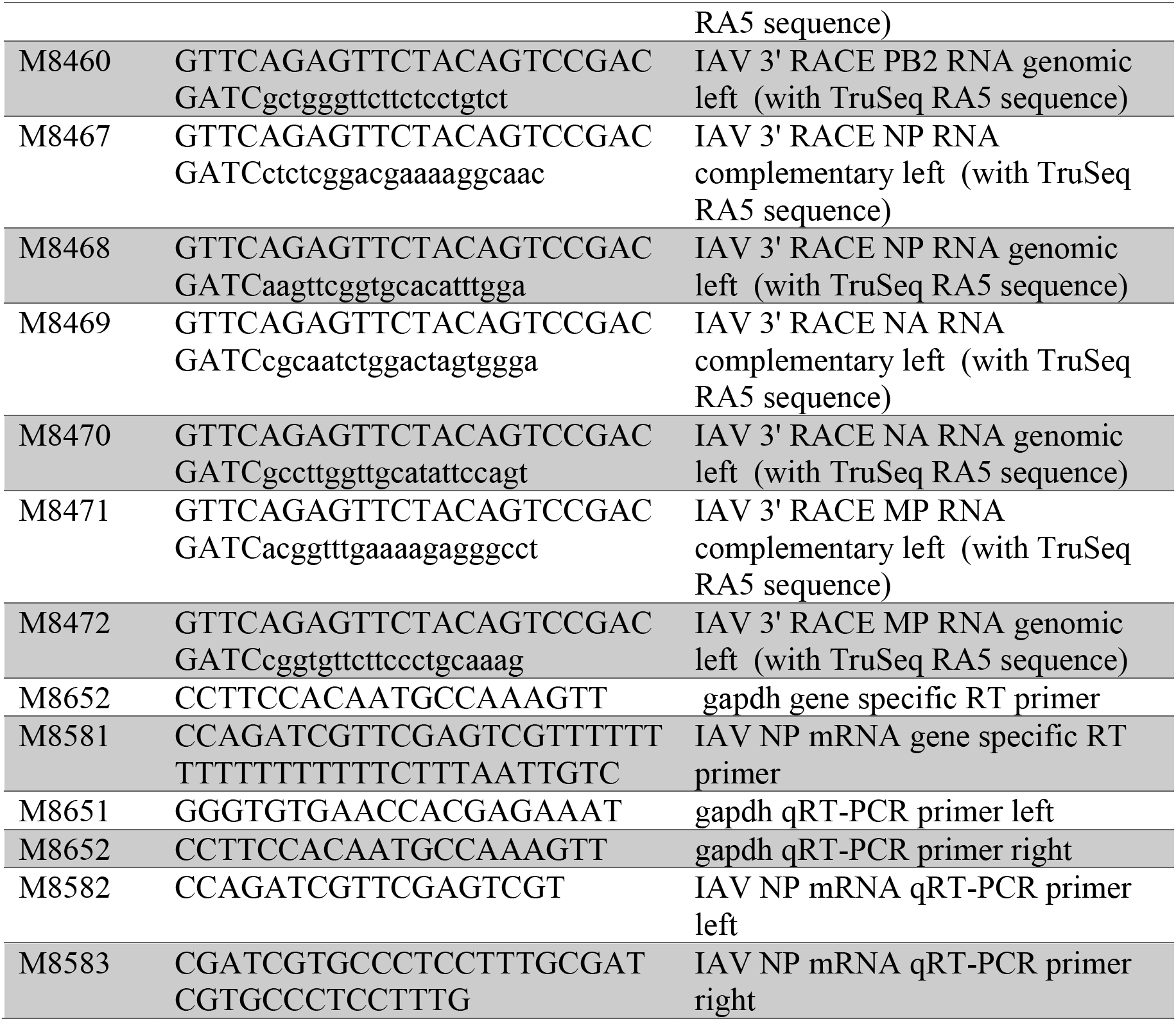
Primers used in this study

